# Mutations in bacterial genes induce unanticipated changes in the relationship between bacterial pathogens in experimental otitis media

**DOI:** 10.1101/464321

**Authors:** Vinal Lakhani, Li Tan, Sayak Mukherjee, William C. L. Stewart, W. Edward Swords, Jayajit Das

## Abstract

Otitis media (OM) is a common polymicrobial infection of the middle ear in children under the age of fifteen years. A widely used experimental strategy to analyze roles of specific phenotypes of bacterial pathogens of OM is to study changes in co-infection kinetics of bacterial populations in animal models when a wild type bacterial strain is replaced by a specific isogenic mutant strain in the co-inoculating mixtures. Since relationships between the OM bacterial pathogens within the host are regulated by many interlinked processes, connecting the changes in the co-infection kinetics to a bacterial phenotype can be challenging. We investigated middle ear co-infections in adult chinchillas (*Chinchilla lanigera*) by two major OM pathogens: nontypeable *Haemophilus influenzae* (NTHi) and *Moraxella catarrhalis* (Mcat), as well as isogenic mutant strains in each bacterial species. We analyzed the infection kinetic data using Lotka-Volterra population dynamics, Maximum Entropy inference, and Akaike Information Criteria (AIC) based model selection. We found that changes in relationships between the bacterial pathogens that were not anticipated in the design of the co-infection experiments involving mutant strains are common and were strong regulators of the co-infecting bacterial populations. The framework developed here allows for a systematic analysis of host-host variations of bacterial populations and small sizes of animal cohorts in co-infection experiments to quantify the role of specific mutant strains in changing the infection kinetics. Our combined approach can be used to analyze the functional footprint of mutant strains in regulating co-infection kinetics in models of experimental OM and other polymicrobial diseases.

## Introduction

Otitis media (OM) is a common polymicrobial bacterial infection of the middle ear in children which is caused by three major bacterial pathogens: nontypeable *Haemophilus influenzae* (NTHi), *Moraxella catarrhalis* (Mcat), and *Streptococcus pneumoniae* (Sp) (1). The relationships among these OM pathogens are both direct and indirect in nature. For example, quorum signals (autoinducer -2 or AI-2) secreted by NTHi help Mcat to form a biofilm and survive in the hostile middle ear environment (2). This interaction represents a direct relationship (or an *active* interaction (3, 4)) between NTHi and Mcat. In another case, NTHi stimulates the host immune response in the middle ear that suppresses the growth of Sp (5, 6); this interaction is an example of an indirect relationship (or a *passive* interaction) between NTHi and Sp. The qualitative (cooperative, competitive or neutral) and quantitative (interaction strength) nature of the *active* and *passive* interactions between the OM bacterial pathogens depend on phenotypes specific to bacterial strains and the host response (4, 7). Mechanistic understanding of how these interactions affect pathogenesis of polymicrobial diseases including OM has been a major research goal for developing vaccine candidates and other therapeutic strategies (8, 9).

A common strategy to evaluate mechanistic roles of specific phenotypes of bacterial OM pathogens *in vivo* has been to co-infect animal models with bacterial pathogens obtained from clinical isolates and then assess changes in infection kinetics by replacing a wild type bacterial strain with a mutant strain (2, 6, 10, 11). The mutant strains are designed to produce a loss or gain of specific bacterial phenotypes of interest. However, because the bacterial phenotypes probed by a mutant strain can be tightly intertwined with the phenotypes of the other bacterial pathogens, via *active* and *passive* interactions, this task could become challenging. For example, a mutant strain of Mcat lacking the ability to receive quorum signal from NTHi conceivably results in a decrease in the co-operative interaction from NTHi to Mcat. However, the same mutation could produce unanticipated changes in other relationships such as change in co-operation/competition of Mcat to NTHi. When these unanticipated changes are *strong* regulators of bacterial populations in co-infection experiments, we are required to revise our mechanistic understanding regarding the role of the specific mutation in influencing co-infection kinetics.

To this end, we address the above challenge by developing a framework that provides an answer to the following question: How is it possible to assess if unanticipated changes in the relationships induced by introducing mutant strains of OM pathogens in co-infection experiments are *strong* or *weak* regulators of the bacterial populations in the experiments? We define loss/gain of phenotype(s) in a specific mutant strain as a *weak* regulator when the interactions between bacterial species in co-infection experiments with the mutant strain are modified according to changes in the phenotype(s) as hypothesized for the mutant strain. The change in the phenotype(s) is defined as a *strong* regulator when additional unanticipated interactions are altered in co-infection experiments with the mutant strain. A more precise and formal definition of the *weak* and *strong* regulators are provided in the Materials and Methods section. The answer to the above question will provide a quantitative way to evaluate the mechanistic role of a bacterial gene in affecting the co-infection kinetics. Our framework combines 1) *in vivo* bacterial load measurements in an animal model, with 2) *in silico* approaches comprised of Lotka-Volterra (LV) population dynamic models (12, 13), Maximum Entropy (MaxEnt) inference (14-17), and, Akaike Information Criterion (AIC) based model selection (18). The animal model we used is a *Chinchilla lanigera* experimental OM model (11), wherein the animals’ middle ears are co-inoculated with NTHi (86-028NP) and Mcat. These strains may be wild type or isogenic mutant strains.

Our study revealed three important findings. First, the unanticipated changes in the relationships between OM bacterial pathogens that substantially affect the co-infection kinetics are commonly present in co-infection with mutant bacterial strains in experimental OM. Second, several bacterial phenotypes are tightly correlated across co-infecting bacterial stains. Third, our combined framework provides a systematic way to deal with two common difficulties faced when analyzing infection kinetic measurements in animal models: host-to-host variations of bacterial populations, and small size of animal cohorts. Our framework can be used to design mutant strains to generate desired infection kinetics in experimental models of polymicrobial diseases such as OM and infections secondary to cystic fibrosis (3) with potential implications for therapeutic design (9).

## Results

### 1. Development of a framework to assess effects of genetic mutation of the bacterial strains in co-infection kinetics

We developed a two species LV model to describe co-infection kinetics of populations of NTHi and Mcat strains within an individual chinchilla host (Fig. 1A and Materials and Methods section). For simplicity in the mathematical expressions of the probability distributions and interaction parameters, we will refer to NTHi strains (wild type or mutant) as species #1 and Mcat strains (wild type or mutant) as species #2 throughout the manuscript. The LV interactions (α_11_, α_12_, α_21_, and, α_22_) characterize the relationships between the bacterial species that originate due to *active* and *passive* interactions (Fig. 1A and Materials and Methods section). Now we can pose the motivating challenge in terms of the LV interactions. How is it possible to assess if unanticipated changes in the LV interactions induced by mutant strains are *strong* or *weak* regulators of the bacterial populations *in vivo*? The unanticipated LV interactions in co-infection experiments with mutant strains are the ones that were not accounted for in the design of the experiment. For example, the *hag* mutant of Mcat does not adhere to the host’s epithelial cell layer as well as the wild type Mcat strain (19). Hence, replacing the wild type Mcat strain in the co-infection by NTHi(*wt*)+Mcat(*wt*) by the *hag* mutant strain should increase Mcat’s self-inhibition; i.e. α_22_ should increase (see Table I for details). Therefore, in the design of the co-infection with NTHi(*wt*)+Mcat(*hag*) one would anticipate a lower carrying capacity for Mcat or an increase of α_22_. However, the changes in *passive* interactions induced by the *hag* mutant can also lead to changes in other LV interactions that were not anticipated, such as an increase in α_21_. In addition, the host-host variations measured in the *NTHi* and *Mcat* populations were assumed to arise from the variations of the LV interactions ({α_ij_}) in our models. We developed a framework (Fig. 1B) to address the above question. The framework is executed in two main steps.

**Figure 1.**
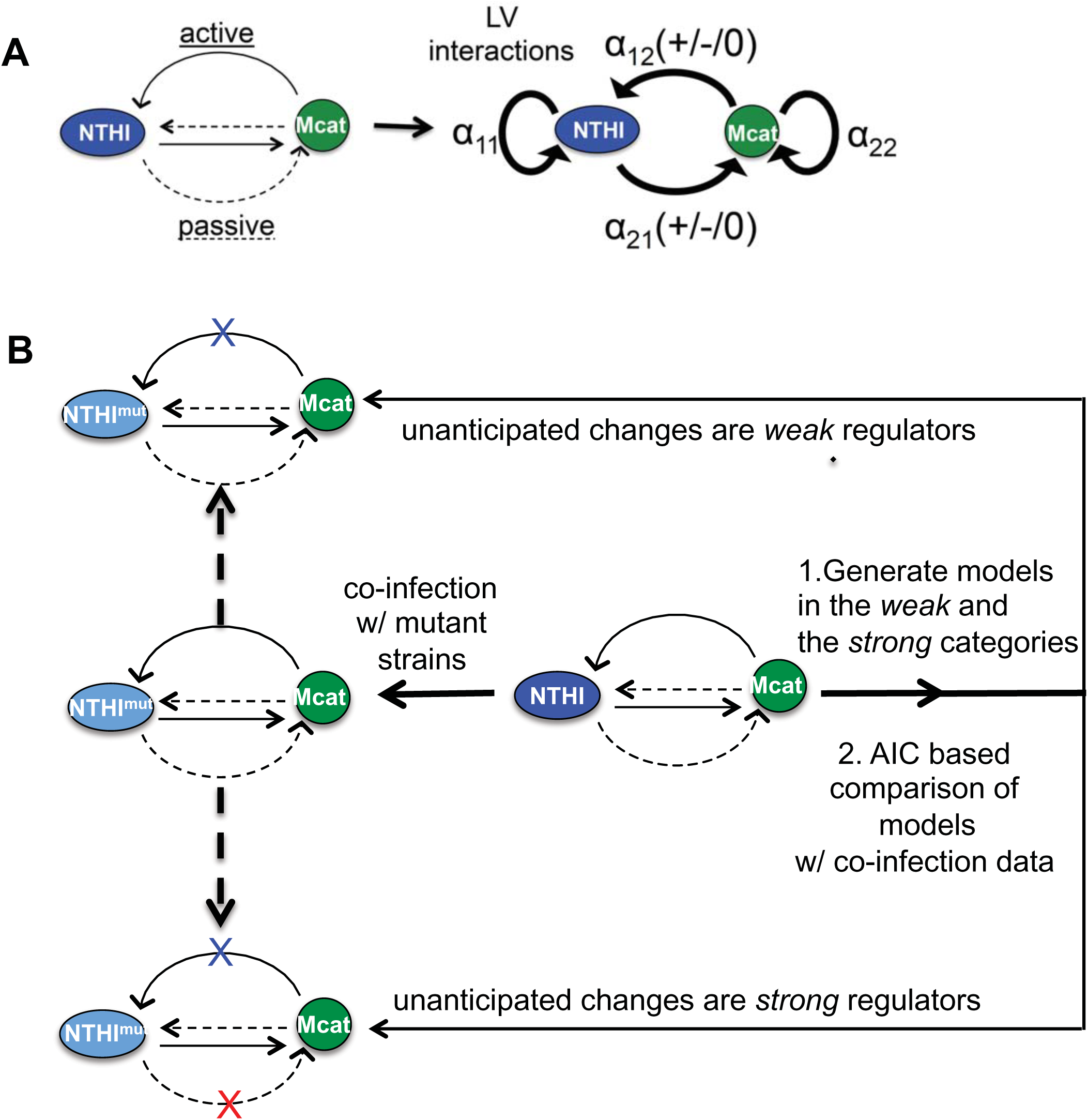
Schematic representation our framework to determine roles of mutant bacterial strains in regulating co-infection kinetics. **(A)** Inter- and intra- species interactions between two bacterial species NTHi and Mcat residing within a host can be both *active* (solid lines) or *passive* (dashed lines) in nature. These interactions can be simplified and described by LV interaction parameters ({α_ij_}). α_11_ (>0) and α_22_ (>0) represent intra- species interactions for NTHi and Mcat, respectively. α_12_ and α_21_ represent the overall effect of Mcat on the growth of NTHi, and, NTHi on the growth of Mcat, respectively. α_12_ and α_21_ can be positive (competitive interaction), zero (neutral interaction), or negative (co-operative interaction). **(B)** Replacing a wild type strain by a mutant strain in the co-infection experiments can change the LV interactions. These changes may be anticipated (blue ‘X’) or not anticipated (red ‘X’) based on the design of the experiment. Our framework uses data from co-infection experiments involving the wild type strains to generate models that determine if these unanticipated changes are *weak* or *strong* regulators of bacterial kinetics. These models are compared to each other using AIC.

**Table I:**
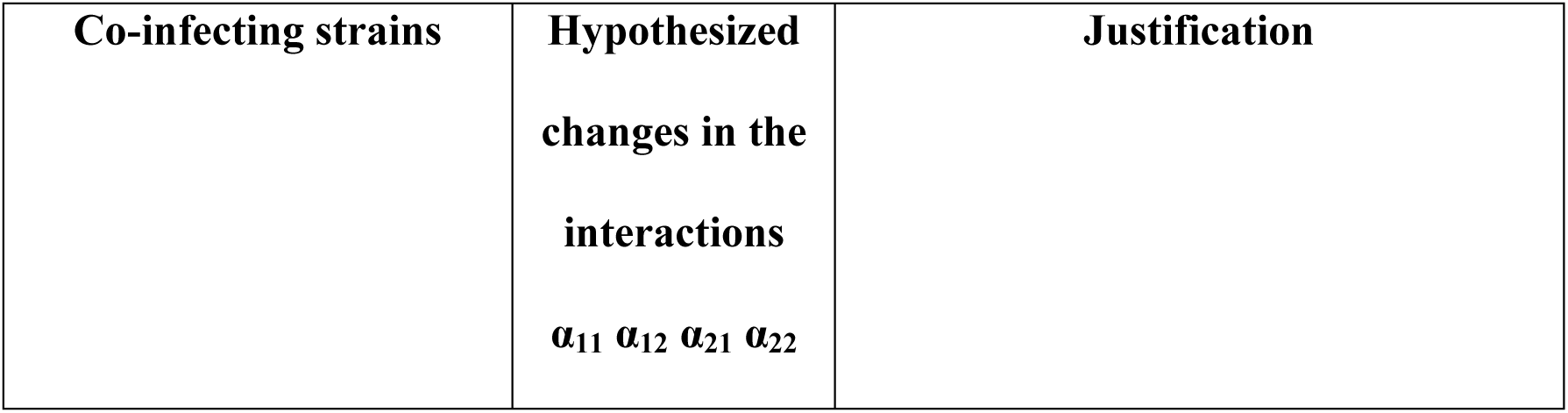
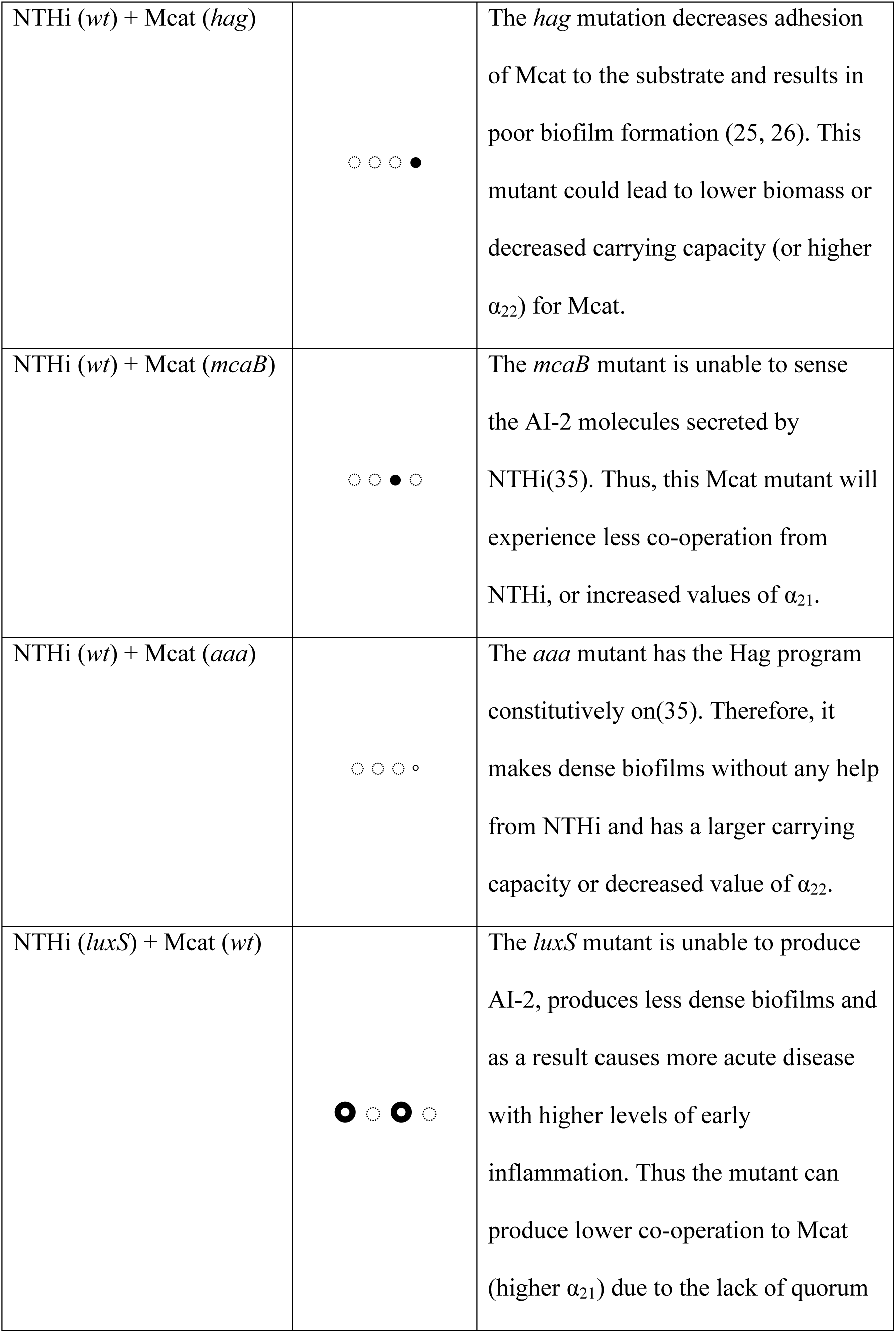
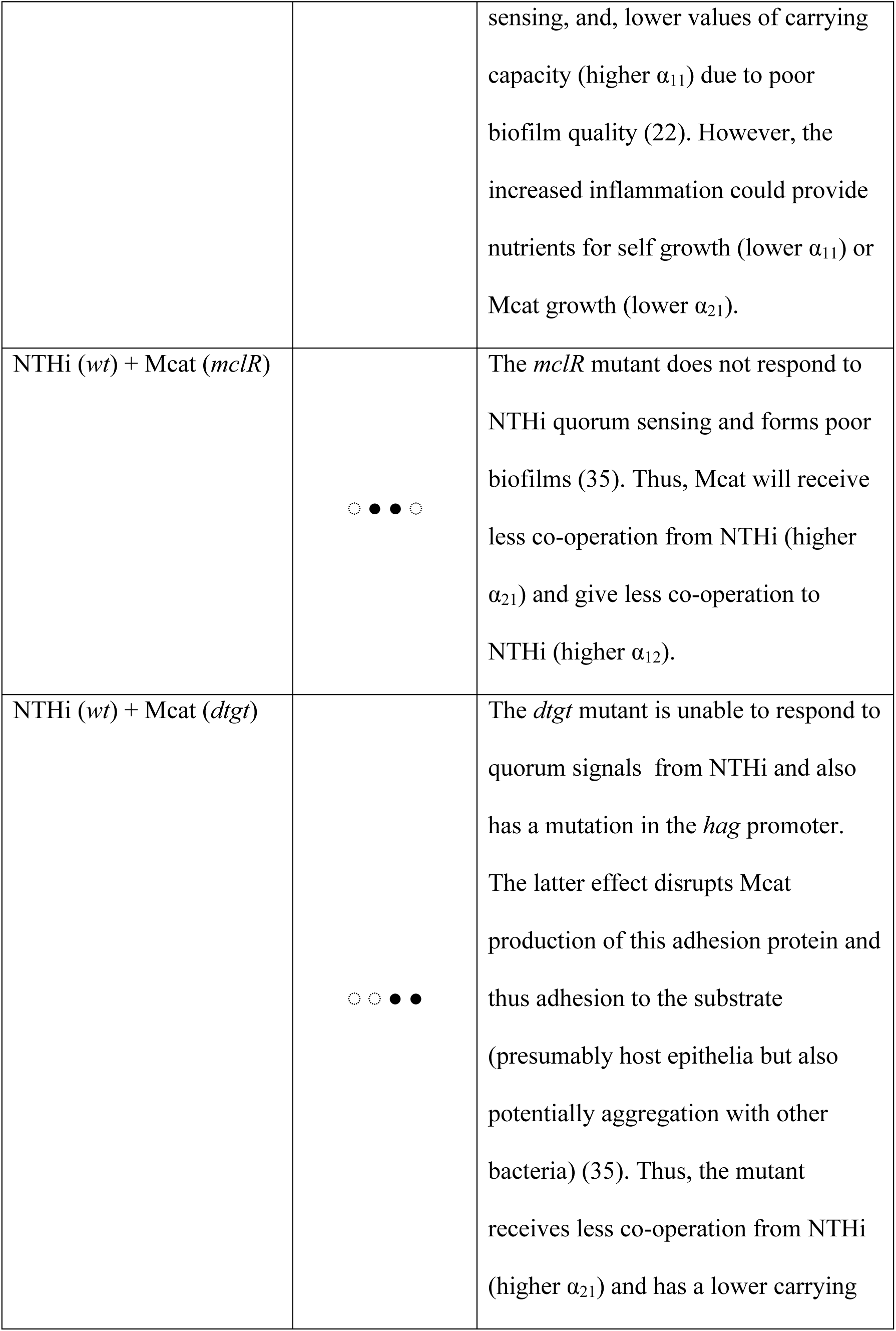
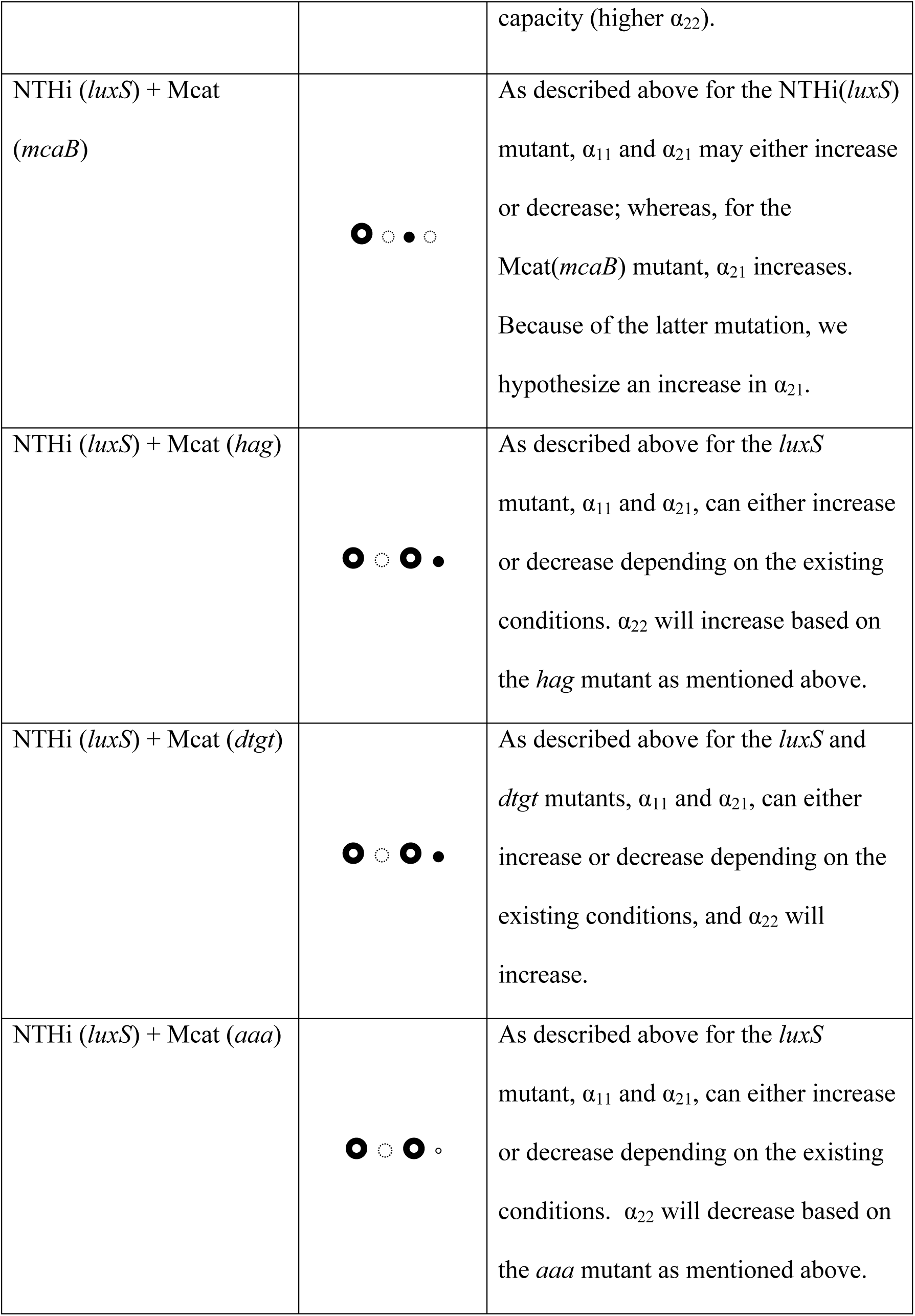
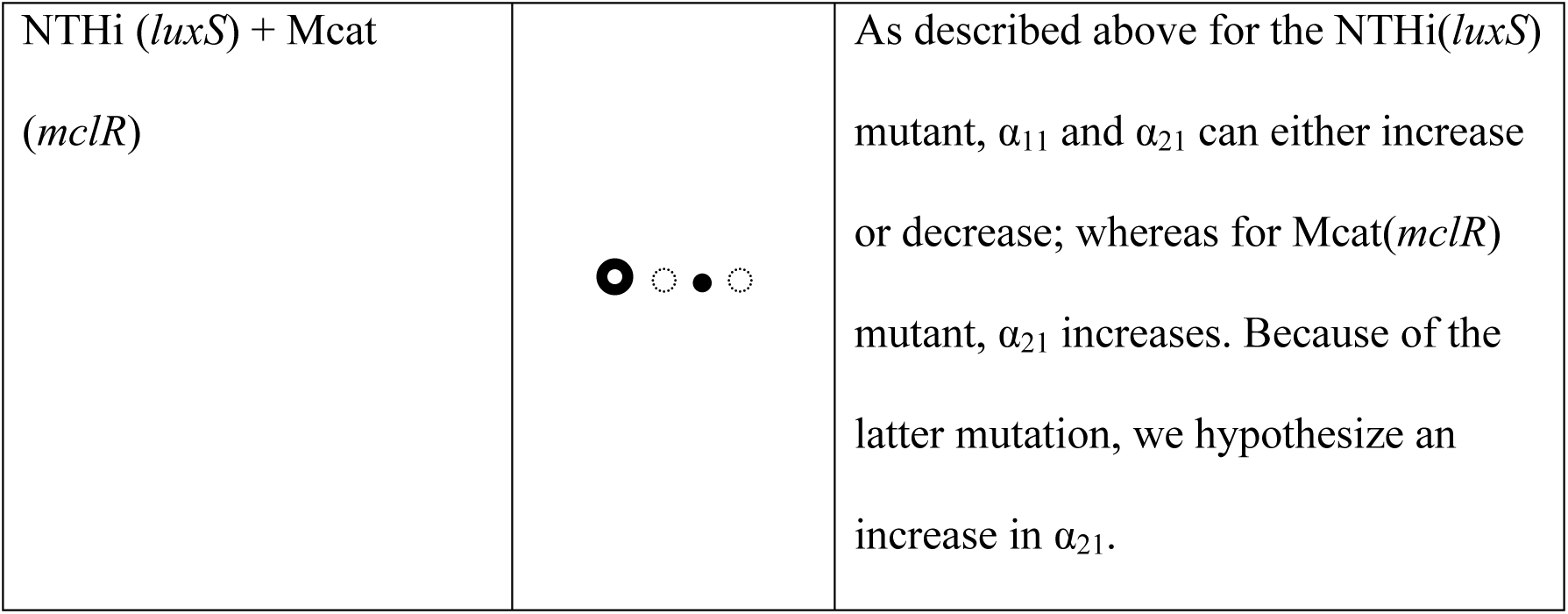
Hypothesized changes in the interactions for co-infections with mutant strains of NTHi and Mcat. For each set of co-infected strains, we indicate the hypothesized changes in the four interaction parameters as suggested by the published literature. An open, dashed circle (◌) indicates no change. A filled circle (●) indicates an increase. A small, open circle (◦) indicates a decrease, and a bull’s-eye 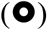 indicates the parameter could increase or decrease. As shown in Fig. 1A, the LV interaction parameters α_11_ and α_22_ represent the strength of NTHi’s and Mcat’s self-inhibition respectively. The α_12_ parameter represents the interaction of Mcat on NTHi, and α_21_ represents the inverse. For the latter two, a decreasing (or more negative) value indicates co-operation; while, an increasing (or more positive) value indicates competition.

#### Step#1

The wild type NTHi (or N_1_) and the wild type Mcat (or N_2_) populations in individual chinchillas were measured in co-infection experiments carried out in a cohort of *n* number of chinchillas at a time *T* (7 or 14 days) post inoculation. These data were used to generate the reference data (denoted by the subscript *r*) *_n_D_r_*. This reference dataset was composed of the mean values 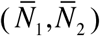, variances 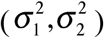, and the covariance (*ρ*_12_) calculated from the bacterial loads measurements in the cohort; *i.e*., 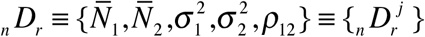 for *j* = 1,…,5. We estimated the probability distribution function 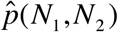 of populations of wild type NTHi (N_1_) and wild type Mcat (N_2_) in the chinchilla cohort using MaxEnt (Fig. S1); wherein, the mean values, variances and the covariance as measured in the experiments (*_n_D_r_*) were constrained in the calculation (Supplemental Material, Section 2). Using this 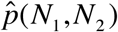, we calculated the joint probability distribution of the interaction parameters 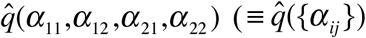 using a MaxEnt based method (details in the Materials and Methods section and Figs. S2-S3). The estimated joint probability distribution function 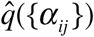 was used to generate models that fall either in the *weak* or the *strong* category.

#### Step#2

We generated a test dataset using data from co-infection experiments where *at least* one of the wild type bacterial strains was replaced by a mutant strain. The test dataset _n′_*D_x_* contains the population means, variances and covariance for the cohort which contained *n′* number of chinchillas for the same time *T* (7 or 14 days) post inoculation. The subscript *x* in _*n*′_*D_x_*, to be determined in our analysis, quantifies the role of the mutation in the co-infection: *x*=*weak* or *x*=*strong*. As described above, this distinction indicates whether the unanticipated changes in LV interactions induced by the mutant strain(s) were *weak* or *strong* regulators of the bacterial populations. In order to safeguard against small sizes of *n′* (~10 or less), we performed bootstrapping (20) on the data; wherein, we sampled *n′* data points with replacement from the original. In this way, we generated *t* sets and determined *x* in each of those *t* samples of _*n*′_*D_x_* These *t* samples of 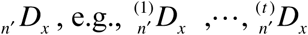 were generated by evaluating _*n*′_*D_x_* in *t* independent groups containing *n′* number of animals each. Next we evaluated which model(s) generated in Step#1 for a particular co-infection experiment best described the *t* samples of the test dataset 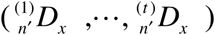; subsequently, we assigned the category (*weak* or *strong*) of the best model to *x*. The best model was found (whenever possible) by comparing the Akaike Information Criterion (AIC) values for each model in a head-to-head pair wise manner for each of the *t* samples. The “Condorcet Winner” was the model which was preferred over all others in head-to-head comparisons(21). We chose the Condorcet winner as the best model. Further details are provided in the Materials and Methods section and in the supplementary material.

### 2. Application of the framework on synthetic data

To test the efficacy of our method, we generated synthetic data and applied the framework developed in the previous section. We had the following three goals in mind: (i) validate the framework, (ii) determine how the strengths of the mutations and/or the host immune response affect interspecies interactions, and (iii) determine the dependence of the model selection on the sample size or the number (*n′*) of animals in the cohort. We generated the synthetic data by numerical solution of coupled ODEs that described LV type population kinetics involving two interacting bacterial species and a host immune response (see Methods and Materials section). The parameters describing the inter- and intra- species bacterial interactions as well as the host immune response were drawn from uniform distribution within specific ranges to generate host-host variations of the co-infection kinetics (see Materials and Methods). We chose the parameter range for the wild type strains such that it produced steady state population values similar to those observed *in vivo* (Figs. S6 – S8).

The average statistical variables 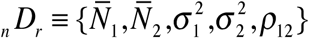, calculated from the numerical solutions at T=day 7, produced the reference dataset. We generated test data sets {_*n*′_*D_x_*} wherein one of the wild type strains was replaced by a mutant strain. Specifically, we considered the following two mutant strains for species #1. The 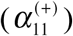 mutant, which is an increase in the α_11_ parameter, possesses increased self-competition for species #1 compared to wild type. The 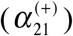 mutant, which is an increase in the α_21_ parameter, possesses increased competition of species #1 towards species #2 as compared to wild type. We also considered two mutant strains for species #2. The 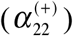 mutant, which is an increase in the α_22_ parameter, possesses increased self-competition for species #2 compared to wild type. The 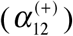 mutant, which is an increase in the α_12_ parameter, possesses increased competition of species #2 towards species #1 as compared to wild type. The mutants were generated by changing the ranges of the associated parameters from that of the wild type strains (see Methods). For example, the range of α_11_ used to generate the mutant strain 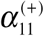 spanned a smaller range [*a′, b*] compared that of the wild type strain, [*a, b*], where, *a*<*a′*. Each of these mutations was performed with low, moderate, and large strengths based on the relative change in the magnitude of range for the parameters. We also solved the co-infection kinetics in the presence of no, weak and strong host immune response. Thus, in total we considered 9×4 different mutation experiments *in silico* (Fig. S9). The test datasets {*_n′_D_x_*} were obtained from the co-infection kinetics involving the above mutant strains.

We show results for two mutant strains 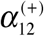 and 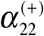 with moderate strength mutations in the absence of any host immune response (Figs. 2B – C). The rest of the mutants are described in the supplementary material. We followed the steps described in section 1 to determine the nature of the mutation (or x) in a test data set *_n′_D_x_*. First, we used *_n_D_r_* to generate models that belonged to the *weak* or the *strong* category corresponding to the co-infections strain#1(wt)+strain#2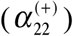 or strain#1(wt)+strain#2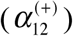. Next, we compared the models with the samples (t>100) of the test dataset and evaluated the Condorcet winner model. We found that for the co-infection with strain#1(wt) + strain#2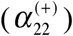, a model in the *x*=*weak* category was the Condorcet winner (Figs. 2D and 2F). In contrast, for the co-infection with strain#1(wt) + strain#2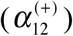, a model in the *x*=*strong* category was the Condorcet winners (Figs. 2E and 2G). These results can be explained by correlations among the LV interaction parameters (Fig. S6) pertaining to the reference data set *_n_D_r_*. The correlations of α_22_ with the other LV parameters for the above co-infection (strain#1(*wt*)+strain#2(*wt*)) is substantially small (<0.06); therefore, increasing α_22_ alone, as in the 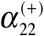 strain, will minimally affect the other parameters. Therefore, the changes in other LV parameters (unanticipated changes) induced by the increase in α_22_ for the 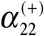 strain will be small. These unanticipated changes would play a *weak* role in regulating the bacterial populations. In contrast, α_12_ is correlated strongly with several other interaction parameters (e.g., α_21_, Corr ≈ -0.5) (Fig. S6). Thus increasing α_12_ even by a moderate amount, as in the 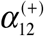 strain, will produce large changes in the other LV parameters. Those unanticipated changes will generate a *strong* effect on the bacterial populations. These expectations about the effects of the correlations among the parameters were consistent with the results obtained from our framework. Therefore, these results validate our framework. The roles of the above mutant strains in affecting co-infection kinetics for different mutation strengths change depending on the mutation strength and/or the presence of the host immune response (Fig. S9). Therefore, strengths of the mutations and the host immune response are important in determining the influence of the mutations in the co-infection kinetics.

**Figure 2.**
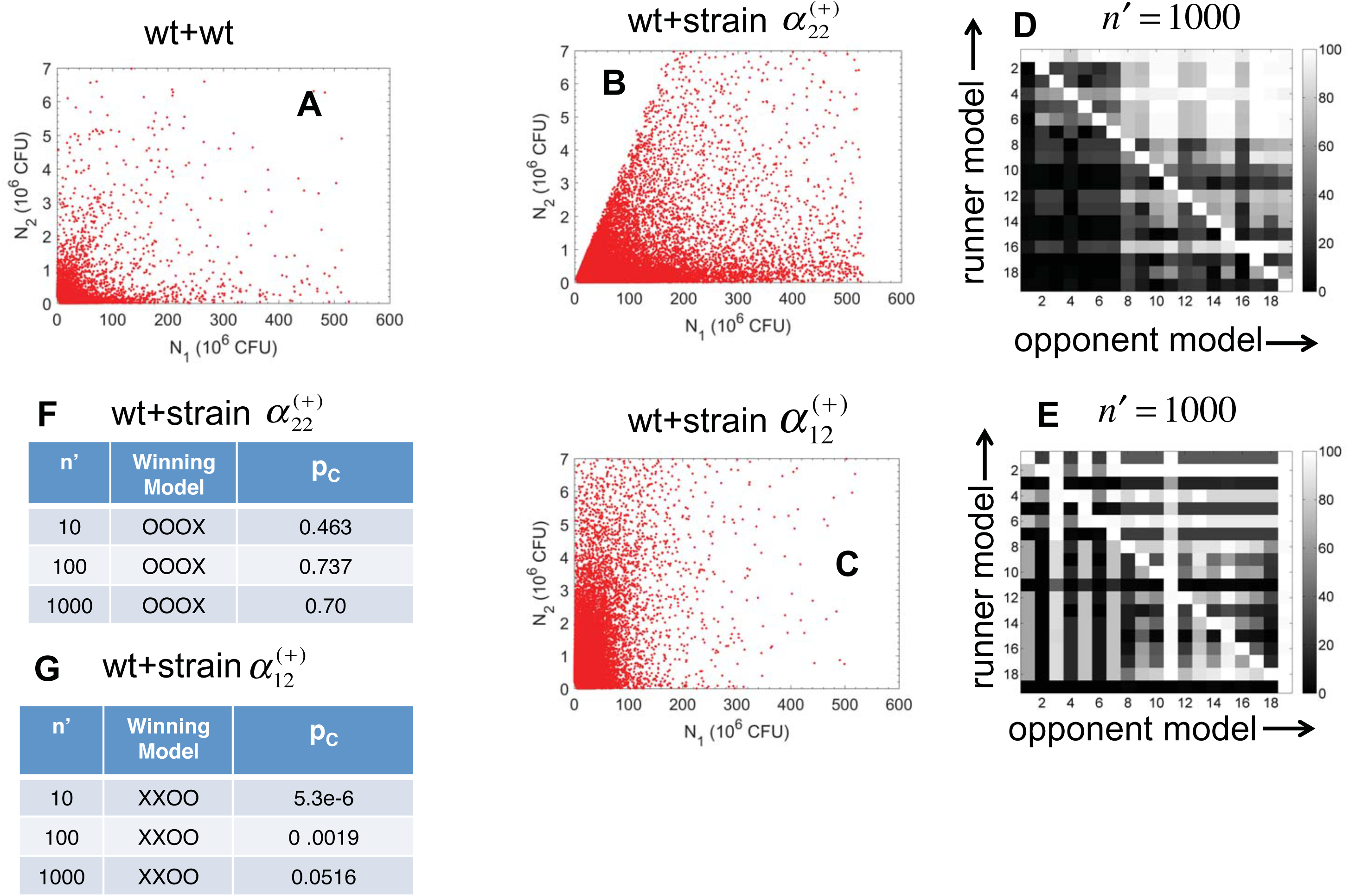
Application of the scheme on synthetic data. **(A)** Shows values of N_1_, N_2_ pairs (10^4^ pairs) obtained from steady state solutions of the ODEs corresponding to the LV model where {α_ij_} were drawn from uniform distributions in the following ranges: 2.74×10^-3^≤ α_11_≤0.2, -200≤α_12_≤5, -5≤ α_21_≤0.1, and, 1.9≤ α_22_≤140. The solutions where either N_1_ or N_2_ went to zero values or became very large (N_1_>530 or N_2_>7) were not included in the synthetic dataset. **(B)** Synthetic data (10^5^ data points) for a co-infection with the mixture wt+α^(+)^_22_ strain. The α^(+)^_22_ strain was generated by increasing the lower range of α_22_ to 120. **(C)** Synthetic data (10^5^ data points) for the co-infection for the mixture wt+α^(+)^_12_ strain. The α^(+)^_12_ strain was generated by increasing the lower range of α_12_ to -2. **(D)** Shows the percentage of the time a runner model won against an opponent model in head-to-head comparison of AICs for the models describing the synthetic data in (C). The m=19 different models are indexed by integers. The percentages shown were obtained for t=100 trials, each with a sample size of n*′*=1000. A bright row indicates the winning model. **(E)** Shows results in head-to-head comparisons between the models presented similar to the data in (D). **(F-G)** The probability p_C_ for the Condorcet winner to win all the pair-wise encounters in the t samples is shown for increasing sample size n*′*. p_C_ for the Condorcet winner model (#i) is calculated using p_C_ = ∏_j(≠i)_ f_ij_, where f_ij_ (>1/2) denotes the fraction of the t samples where the Condorcet winner model #i was preferred over model #j. The product is calculated for all the m-1 pair wise combinations where m number of models were considered. p_C_ increased with the sample size (n*′*).

We checked the dependence of the Condorcet winner on the sample size *n′* in the test dataset. We found that even for small sample sizes (*n′*=10), the framework picked the correct Condorcet winner, however, the margin of victory increased with larger *n′* (Figs. 2F and 2G).

### 3. Analysis of the *in vivo* data

We analyzed co-infection kinetics in *Chinchilla lanigera* co-inoculated with a mixture of wild type NTHi and Mcat strains or mixtures of NTHi and Mcat strains wherein at least one of the bacterial strains was a mutant strain. The chinchillas were inoculated by injecting 10^3^ CFU of NTHi and 10^4^ CFU of Mcat directly into the middle ears of the animals. The co-infection experiments investigated the *hag, mcaB, aaa, mclR*, and *dtgt* mutant strains of Mcat and the *luxS* mutant strain of NTHi (Table I and S1). The properties of the mutant strains and their hypothesized effects on the LV interaction parameters are described in Table I. The populations of the NTHi and Mcat strains were measured at 7 days (day 7) and 14 days (day 14) post inoculation (Fig. 3 and Table S1).

**Figure 3.**
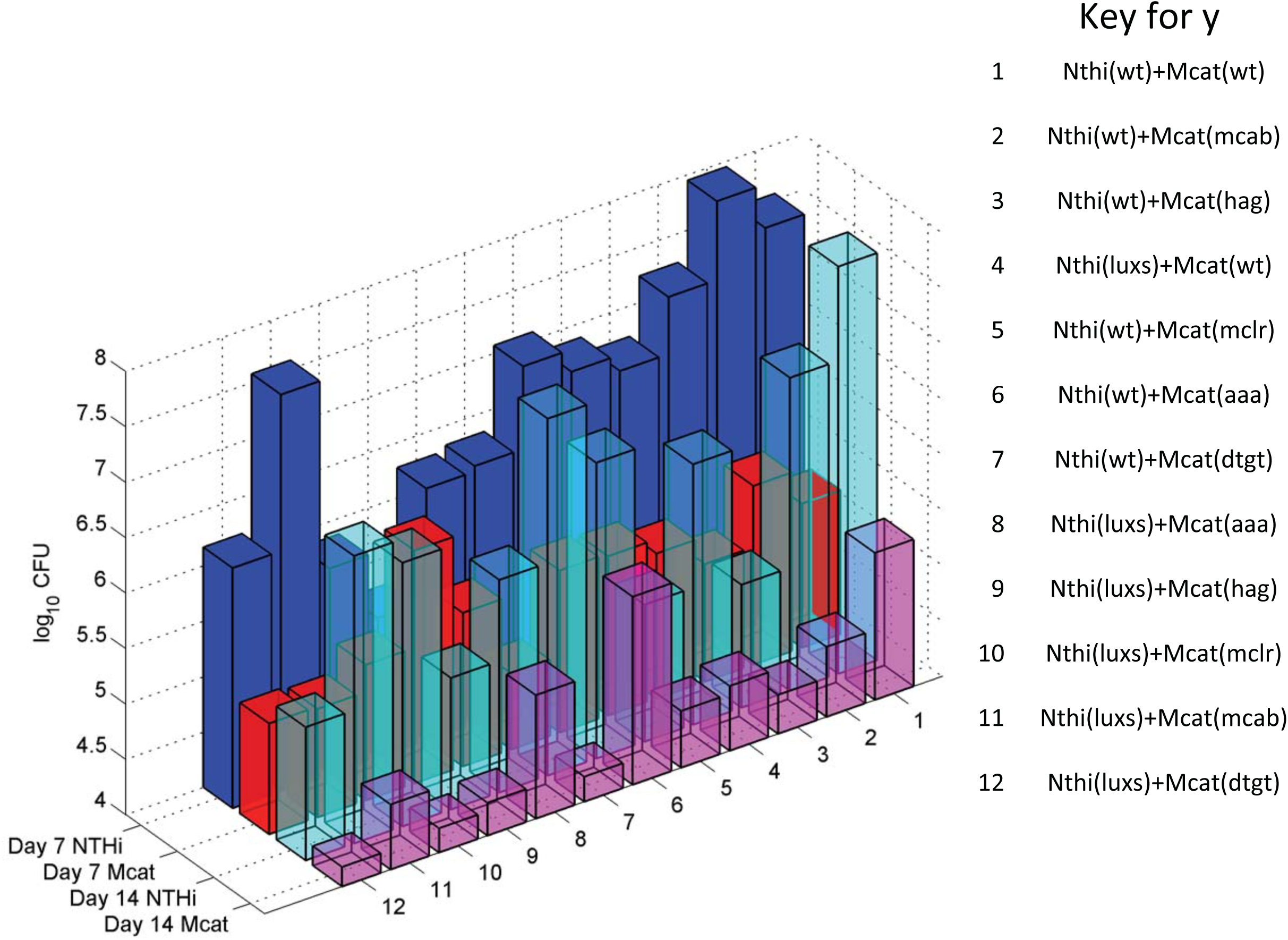
Mean populations of NTHi and Mcat strains in co-infection experiments. Shows the mean populations of NTHi and Mcat strains at day 7 and day 14 post inoculation calculated from counts in the bullae measured in >5 chinchillas for each of the 12 different cases of co-inoculation. The combinations of the bacterial strains used for co-inoculating the chinchillas are listed.

The data were collected from both ears of the chinchillas in cohorts containing more than five animals. The bacterial counts showed large to moderate host-to-host variations (Fig. S1). These variations could arise from the host-host differences in the physiology, anatomy, and immune responses in the upper respiratory tract of the outbred population of chinchillas. The mean NTHi population was substantially larger (>100 fold) than that of Mcat for the co-inoculation with the wild type NTHi and Mcat strains (Fig. 3). The mean populations of the wild type NTHi strain at day 7 was lower if co-inoculated with any mutant Mcat strain (except Mcat(*mcaB*)) rather than if co-inoculated with wild type Mcat (Table S1). In contrast, the mean populations of the Mcat strains for the same co-infections showed small changes (increase or decrease) (Fig. 3 and Table S1). Coinoculation with NTHi(*luxS*)+Mcat(*wt*) resulted in a negligible change in the mean Mcat population but a large decrease in the mean NTHi population at day 7 as compared to the NTHi(*wt*)+Mcat(*wt*) experiment. When both the NTHi(*wt*) and the Mcat(*wt*) strains were replaced by their mutant strains in the co-inoculating mixtures the mean populations of both the strains decreased at day 7. At day 14, the mean populations of the NTHi strains decreased, compared to the NTHi(*wt*)+Mcat(*wt*) experiment, in all the experiments with any mutant strains. The covariances between the populations of the NTHi and the Mcat strains were negative for the majority of the cases investigated here (Table S1). In a few cases, such as NTHi(*wt*) +Mcat(*hag*) NTHi or NTHi(*wt*)+Mcat(*mcab*) the covariances were positive (Table S1). Overall, the data showed a complex pattern as further explained below.

Changes in the bacterial counts due to mutations pointed towards the presence of unanticipated changes in the bacterial relationships in regulating bacterial populations. For example, compared to co-infecting with NTHi(*wt*)+Mcat(*wt*), co-infecting with NTHi(*wt*)+Mcat(*hag*) substantially decreases the NTHi population (almost by half) whereas the Mcat population only decreases a small amount. Since the *hag* mutant has lower adherence and poor biofilm formation capability compared to its wild type counterpart, we would expect the Mcat population to decrease while having minimal consequence to the NTHi population. Because the NTHi population was substantially reduced here, it suggests a potential change in the interaction from Mcat towards NTHi. We used our framework to quantify the roles of specific NTHi and Mcat mutations in regulating the bacterial populations in co-infection experiments. As described in the previous section, the data from the co-infection experiments with the wild type strains generated our reference dataset *_n_D_r_*. The models in the *weak* or the *strong* category were generated using *_n_D_r_* and were compared against the test datasets {*_n′_D_x_*}. The test datasets were obtained from the co-infection experiments that involved *at least* one mutant strain. Different samples of a test data set were obtained by using bootstrapping (20). Our analysis showed that for the majority of the cases, models with additional interactions (*strong* models) better described the data (at both day 7 and day 14) compared to models with no additional interactions (*weak* models) (Fig. 4). We found that the *weak* model described the data obtained at day 7 for the co-infection with NTHi (*wt*)+Mcat (*mcaB*) (Fig. 4B) better than any *strong* model. However, at day 14, a *strong* model (Fig. S4) described the same data better than any *weak* model. Therefore, unanticipated changes in LV interactions were prevalent in co-infections with mutant bacterial strains both at early and late stages of the co-infection kinetics.

**Figure 4.**
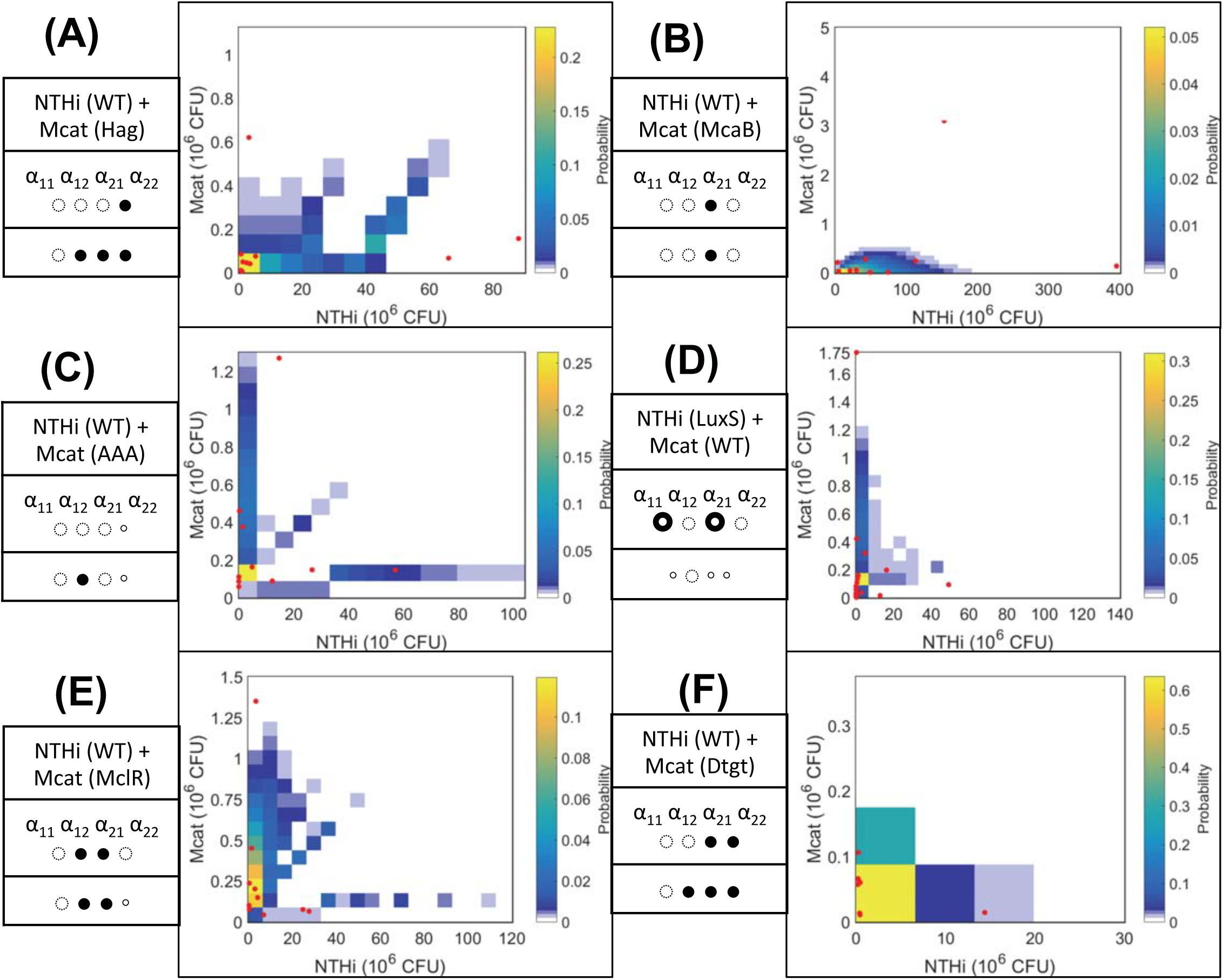
Comparison between the Condorcet winner model and measurement at day 7 post inoculation with mutant strains. The probability distribution function 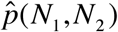 generated by the Condorcet winner model for a co-infection at day 7 post inoculation is shown using a heat map. The measured bacterial loads for the same co-infection for individual chinchillas are shown in red points. The anticipated changes in the LV parameters for a co-infection involving a specific mutant strain are shown in the first row of the table shown on the left of a sub figure. The changes suggested by the Condorcet winning model are shown in the second row. A filled (●) or a smaller empty circle (◦) indicates an increase or decrease of a specific parameter, respectively. The cases where the phenotype is uncertain, i.e. either an increase or decrease, are marked by a bullseye symbol 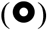. **(A)** NTHi (wt)-Mcat (*hag*). **(B)** NTHi (wt)-Mcat (*mcaB*). **(C)** NTHi (wt)-Mcat (*aaa*). **(D)** NTHi (*luxs*)-Mcat (WT). **(E)** NTHi (wt)-Mcat (*mclR*). **(F)** NTHi (wt)-Mcat (*dtgt*). Note that we compared our prediction against the measured data in terms of average populations as bacterial measurements were available for only few animals. The individual data points shown on the graphs were not explicitly compared, thus, some of the individual measurements (dots) can lie at the boundaries of the predicted distribution (colored squares) and need not reflect quality of comparison between the average bacterial populations.

### 4. Host immune responses modulate Mcat-NTHi interactions at later stages of the infection

The mean populations of the wild type strains of NTHi and Mcat increase as the infection progresses from day 7 to day 14 post inoculation (Fig. 5). However, the covariance of the NTHi and Mcat populations becomes more negative (~2 fold change) (Table S1). The negative correlation indicates the populations of the two species are more mutually exclusive; that is, when one species has high abundance, then the other’s is low. We hypothesized that the host immune response generated by both the pathogens could lead to a decrease in co-operation (or increase in α_12_ and α_21_) between Mcat and NTHi. We tested our hypothesis by applying our scheme with the day 7 data as the reference and day 14 data as the test. The model wherein α_12_ and α_21_ increase was the Condorcet winner (Fig. 5). The agreement demonstrates the role of the host immune response in regulating *passive* inter-species interaction between Mcat and NTHi.

**Figure 5.**
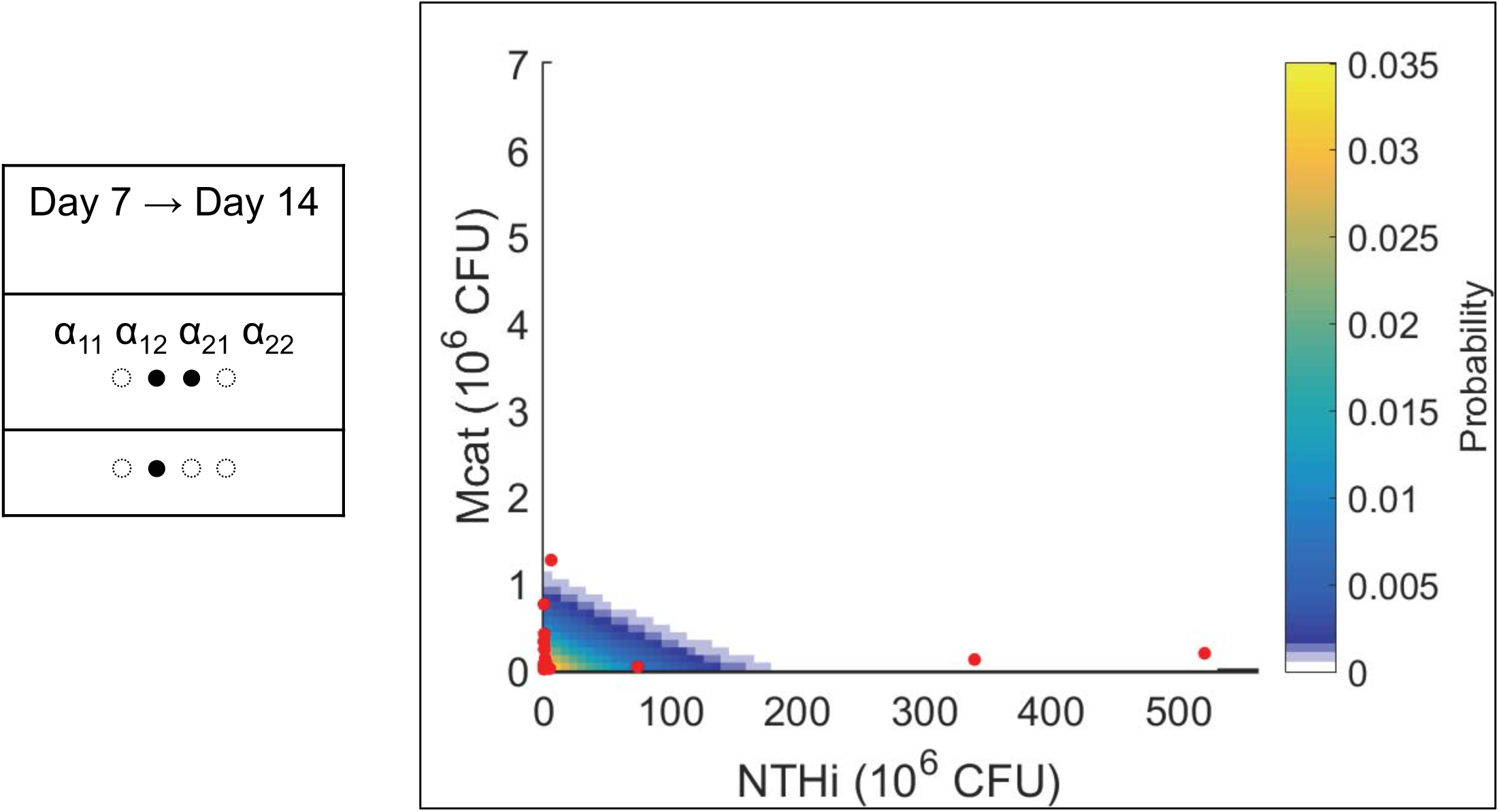
The prediction at day 14 post inoculation for the wild type strains generated using the day 7 data. The data are displayed using the same visualization scheme as in Fig. 4.

## Discussion

Co-infection of animal models with mutant bacterial strains is a powerful tool in probing mechanisms that underlie pathogenesis of polymicrobial infections such as OM. However, the interconnected and variable nature of interactions involving bacterial pathogens within the host makes it challenging to connect specific perturbations, such as a mutation, in these experiments to mechanisms. The data-driven framework developed here provides a systematic method of addressing this challenge. The framework uses bacterial counts measured in animal hosts to quantitatively determine perturbations in the bacterial interactions induced by the replacement of a wild type bacterial strain with a mutant strain that strongly regulates bacterial populations in the co-infection. Therefore, using this framework we are able to quantitatively assess the mechanistic role a specific bacterial phenotype probed by an isogenic mutant strain in affecting the co-infection kinetics. Isogenic mutant strains are used for identification of bacterial determinants of colonization, persistence and virulence. Thus the quantitative information obtained from our framework will be valuable for determining specific targets for diagnostics, development of therapy and potentially vaccination. In addition, our framework also addresses the practical problem of systematically analyzing bacterial count data obtained from a small size (~ 10 animals) of animal cohorts in co-infection experiments.

Application of our framework to co-infection experiments in *Chinchilla lanigera* coinoculated with wild type and mutant strains of two major OM pathogens (NTHi and Mcat) found that in a majority of the co-infections the mutant strains gave rise to unanticipated changes in the bacterial interactions, which influenced the bacterial populations substantially. The emergence of unanticipated perturbations of the bacterial interactions is likely caused by the interdependencies between the interactions as well as the hostile environment of the host (16). The interdependencies can be caused due to many shared processes such as feeding on common nutrients, exchanging small molecules, and the host immune response that regulates the growth of OM pathogens within the host (5, 8). Therefore, when a specific bacterial phenotype is altered in the form of a mutant strain, several other phenotypes in co-infecting OM pathogens are also altered, some of which can be non-intuitive. Our analysis of the synthetic co-infection data lends support to this speculation. We found that correlations between the LV parameters generated non-intuitive changes in several LV interactions when a specific LV interaction was perturbed in a mutant strain, and, in many cases these unanticipated changes in the LV interactions were *strong* regulators of the bacterial populations.

Our framework required estimation of LV interactions involving the co-infecting bacterial pathogens using the measured bacterial counts. The estimated interactions between wild type strains of NTHi and Mcat demonstrate prevalence of co-operative interspecies interactions (α_12_<0, α_21_<0). Previous experiments noted several molecular mechanisms regarding the help of NTHi towards Mcat’s growth, e.g., the quorum signal AI-2 secreted by NTHi helping Mcat to form a biofilm and thereby helping it to survive within the host (22, 23). Reciprocally, the estimated interactions also suggested cooperation of Mcat towards the growth of NTHi (or α_21_<0). Such a co-operative effect can potentially occur through *passive* interactions. For example, Mcat binds and sequesters AI-2 molecules secreted by NTHi; this sequestration could help keep the AI-2 abundances at an optimal level for production of quorum signals by NTHi. Similar optimal regulation of quorum sensing has been found in mutualistic relationship between two human oral bacteria, *Actinomyces naeslundii* T14V and *Streptococcus oralis* 34, where, AI-2 secreted from the later bacterial species help the former species to form biofilms (24). Furthermore, Mcat bacteria are known to form large aggregates (or autoagglutination) via the Hag protein. Such aggregates can help NTHi to form biofilms as mixed NTHi-Mcat biofilms within the host (25, 26). Another example of cooperation involves nutrient recruitment. The inflammation caused by Mcat can produce an influx of host serum which also can provide nutrients for the growth of NTHi (27). All three of the above sources of interaction could contribute towards generating an overall co-operative interaction from Mcat to NTHi.

We found that several LV interactions (e.g., α_12_ and α_21_) involving NTHi and Mcat are tightly correlated (|Corr|>0.5) with each other. Since the MaxEnt method estimates the most spread out or uniform probability distribution phenotype that is consistent with the measured data (17), the method used here provided the most conservative estimate correlations between the LV interactions. This approach is a major departure from several methods that have been developed in the recent years to evaluate interactions(28-30) pertaining to microbiome datasets where the correlations between the interaction parameters are not analyzed. Ecological models often assume LV interactions between co-existing species as uncorrelated random variables (13). This assumption makes the calculations amenable to analytical methods. The presence of strong correlations between LV interaction parameters could have important implications in assessing general principles underlying the diversity of eco-systems (31).

## Materials and Methods

### A. Experiments

*M. catarrhalis* persistence in the middle ear chambers of chinchillas was assessed essentially as described previously (2). Animals were purchased on need and allowed to acclimate to the vivarium for > 10 days before infection. No used animals showed visible sign of illness prior to infection. Chinchillas (five animals per group) were anesthetized with isofluorane and infected via transbullar injection with both ~ 10^4^ CFU of *M. catarrhalis* and ~ 10^3^ CFU of *H. influenzae*. All inocula were confirmed by plate counting. Animals were euthanized at 7 and 14 days post-infection, and their bullae were aseptically opened to recover possible effusion fluid. Middle ear lavage was performed using sterile PBS and also saved and combined. Bullae were then excised and homogenized in 10 ml of sterile PBS. All of fluid and homogenized samples were serially diluted and plated on brain heart infusion (BHI) agar plates to obtain viable counts of *M. catarrhalis*. Note that *H. influenzae* would not grow on BHI plate due to lacking of hemin and NAD. *H. influenzae* bacteria were enumerated on BHI agar supplemented with 10 micrograms/ml heme and NAD and containing 5 micrograms/ml clarithromycin, which inhibits growth of *M. catarrhalis*.

#### Bacterial strains and growth conditions

*M. catarrhalis* strain O35E, as a WT and parent strain in this study, is a commonly-used laboratory strain (32). *M. catarrhalis* O35E *hag*::Sp containing a spectinomycin resistance cassette disrupting the *hag* gene is a kind gift from Dr. Eric Hansen. Nontypeable *H. influenzae* strain 86-028NP is a nasopharyngeal isolate from a child with chronic OM (33), and its *luxS*::Kn mutant with a kanamycin resistance cassette disrupting the AI-2 synthase was described previously (22). *M. catarrhalis* strains were cultured in BHI medium (Difco), and *H. influenzae* strains were cultivated in BHI medium supplemented with 10 μg/ml of hemin chloride (MP Biomedicals) and 10 μg/ml of NAD (Sigma), referred as supplemented BHI (sBHI). Mixture cultures of both *M. catarrhalis* and *H. influenzae* used sBHI.

#### Mutant bacterial strains

*M. catarrhalis mclR*::spec was generated by insertional mutagenesis using the following approach. Genomic DNA was purified from *M. catarrhalis* O35E using the Wizard genomic DNA purification kit (Promega), essentially according to the manufacturer’s instructions. Portions of the *mclR* (allele MCR_1062) open reading frame were amplified using the PCR and primers specific for intragenic regions (luxRUPF: CATCATGACTTGGTAACTTGCTG, luxRUPR: GCTGATCGGCAATTTGCCCCCGGGGTCGAGTGGCTTCTACACC). The resulting amplicons were cloned and a spectinomycin resistance cassette was introduced into a SmaI restriction site within the intergenic primers. This mutant allele was introduced into the parental strain by natural transformation and the resistant derivatives were confirmed by PCR and DNA sequence analysis.

#### Hag promoter mutants

Deletion of a tgt sequence and insertion of a *aaa* sequence within a predicted lux box within the hag promoter was achieved by overlap PCR; the resulting allele was introduced by natural transformation using a linked spectinomycin resistance marker (Li Tan et al., *under revision*, 2018).

### B. Estimation of probability distribution function q({α_ij_}) of LV interactions {α_ij_}

#### (1) LV modeling of the population kinetics

We described the population kinetics of populations of NTHi and Mcat in the middle ear of the host (*Chinchilla lanigera*) using two coupled Ordinary Differential Equations (ODEs) following the Lotka-Volterra (LV) model (12):

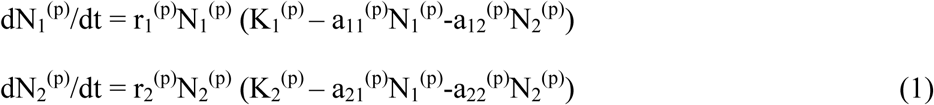

N_1_^(p)^ and N_2_^(p)^ denote the populations of the NTHi and Mcat strains, respectively, in the middle ear of an individual host. Individual hosts are indexed by the superscript p. The parameters r_1_K_1_ and r_2_K_2_ denote the growth rates of the species 1 and 2, respectively, where K_1_ and K_2_ denote the corresponding carrying capacities. The parameters {a_ij_} describe effective interactions involving the bacterial species. a_11_ and a_22_ denote self-competition for growth for species 1 and 2, respectively, and a_12_ and a_21_ denote the influence (competition or co-operation) of species 2 and 1 on the growth of the other species, 1 and 2, respectively. Note, a_12_ and a_21_ can assume positive (indicating competition) or negative (indicating co-operation) values; whereas, a_11_ and a_22_ can only possess non-negative values. The carrying capacities and the interaction parameters {a_ij_} determine the maximum bacterial load that can be sustained in the local environment. The above simple description of the bacterial infection kinetics within a host provides a coarse-grained and effective description of the kinetics, where, the bacterial populations represent an average over spatial length scales including spatial structures such as biofilms. The effect of the immune response, nutrients, protective effects of biofilm formation, and, quorum sensing are effectively described in terms of the interaction parameters and the carrying capacities (Table I). The above model describes the interactions between the bacterial species and the host minimally where the LV interaction parameters provide a clear description of the inter- and intra- species interactions between the co-infecting bacterial species. These effects can vary from host-to-host giving rise to host dependent values of the effective parameters, therefore, we consider host-host variations of the model parameters ({a_ij_}, {K_i_}). Since, the NTHi and Mcat replication rates are ~1 hour^-1^, it is reasonable to assume that in a time scale of days the kinetics in Eq. (1) reaches a steady state, *i.e*.,

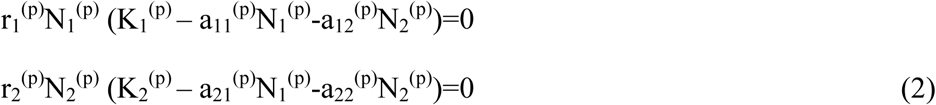

The steady state equations (Eq. 2) help reduce the number of parameters in determining populations of NTHi and Mcat.

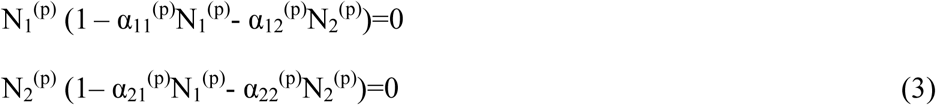

We defined α_ij_^(p)^=a_ij_^(p)^/K_i_^(p)^ in Eq. (3). Thus, the dependences of the carrying capacities {K_i_} are effectively contained in the scaled variables {α_ij_}. The <u>*LV interaction parameters*</u> α_11_^(p)^, α_12_^(p)^, α_21_^(p)^, α_22_^(p)^ determine the bacterial abundances at the steady state (or the stable fixed points). The above equations produce four fixed points, (N_1_ ^(p)^=0, N_2_ ^(p)^=0), (N_1_ ^(p)^=0, N_2_ ^(p)^=1/α_22_ ^(p)^), (N_1_ ^(p)^=1/α_11_ ^(p)^, N_2_ ^(p)^=0), and, (N_1_ ^(p)^=([**α**^(p)^]^-1^)_11_+([**α**^(p)^]^-1^)_12_, N_2_ ^(p)^=([**α**^(p)^]^-1^)_21_+([**α**^(p)^]^-1^)_22_), where, **α**^(p)^ and [**α**^(p)^]^-1^ denote the matrix, {α_ij_ ^(p)^}, and its inverse, respectively. The stability of the fixed points is determined by the linear stability analysis (Section 3, Supplementary Material). We consider only the stable fixed points, where, N_1_ ^(p)^>0 and N_2_ ^(p)^>0. The parameter values yielding any other type of solutions (e.g., N_1_ ^(p)^= N_2_ ^(p)^=0, N_1_ ^(p)^<0 or N_2_ ^(p)^<0, N_1_ ^(p)^ → ∞or N_2_ ^(p)^ → ∞) are considered not to occur in the bacterial kinetics. Thus, we consider the solutions,

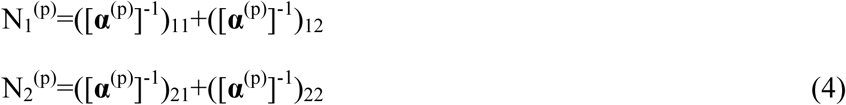

when they are stable and are positive. Next, we estimated the LV interaction parameters {α^(p)^_ij_} from the bacterial loads (N_1_^(p)^ and N_2_^(p)^) for an individual animal (indexed by p). For a measured value of N_1_^(p)^ and N_2_^(p)^, it is not possible to estimate the four parameters {α_ij_^(p)^} uniquely using the above equations. Therefore, we developed a MaxEnt based inference scheme to estimate the parameters in the chinchilla population using the measured bacterial loads. The MaxEnt based method estimates parameters based on the measured data without any additional prior assumption. This also implies that MaxEnt estimates the ‘flattest’ distribution that is consistent with the measured data. A recent work(34) used Maximum Caliber inference, which is an extension of MaxEnt for analyzing time dependent data(15, 17), to estimate parameters in a gene regulatory reaction network. Parameter estimation in gene regulatory reaction networks using sparse time dependent data represents a problem of similar spirit as the problem investigated here. First we estimated the probability distribution function of N_1_ and N_2_, 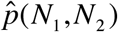, in the chinchilla population using MaxEnt (Fig. S1, Supplementary Material). Then we estimated the joint probability distribution function 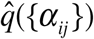 in the interaction parameters {α_ij_} using the estimated 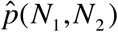 by applying MaxEnt the second time (Fig. S2 – S3). The details regarding the implementation of the method for the *in vivo* data are provided below and in the supplementary material (Section 2, Supplementary Material).

#### (2) MaxEnt estimation of 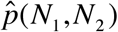

The bacterial loads for wild type strains of NTHi and Mcat in 10 adult chinchillas (10×2 ears = 20 samples) were used to calculated the mean bacterial loads (E(N_1_) and E(N_2_), variances (σ^2^(N_1_) and σ^2^(N_2_)), and the covariance, Cov(N_1_, N_2_). 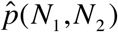 was estimated using a MaxEnt procedure where the space in N_1_ and N_2_ was discretized on an equally spaced 81×81 lattice ({I,J}). The ranges of N_1_ and N_2_ were 0 – 530 × 10^6^ CFUs and 0 – 7 × 10^6^ CFUs respectively. The MaxEnt method involved maximizing Shannon Entropy(16, 17), 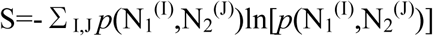 subject to the constraints, 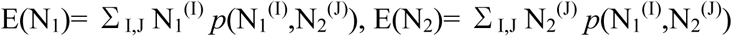, 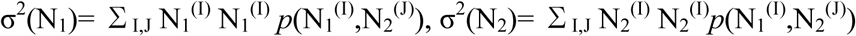, and, 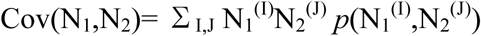. The solution that maximizes S is given by 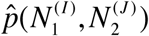 ∝exp(λ_1_N_1_^(I)^+ λ_2_N_2_^(J)^+ λ_3_N_1_^(I)^N_1_^(I)^ + λ_4_N_2_^(J)^N_2_^(J)^ + λ_5_N_1_^(I)^N_2_^(J)^). The five Lagrange’s multipliers (λ_1_,…,λ_5_) were calculated by solving the constraint equations in MATLAB using the built-in function *fsolve*. We will denote the discrete probability distribution, 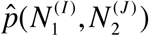 by 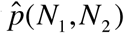, hereafter, to keep the notation simple.

#### (3) Estimation of 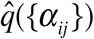

We estimated 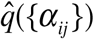 using 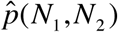 by applying the MaxEnt inference the second time. We discretize the four dimensional space spanned by α_11_, α_12_, α_21_, and α_22_ on a grid. α_11_ and α_22_ assume only positive real values, and α_21_ and α_12_ can assume both positive and negative real values. Specifically, we discretized α_11_ from 0.027 – 2 × 10^6^ CFU and α_22_ from 18.9189 – 1400 × 10^6^ CFU into 74 bins each. We discretized α_12_ from -2000 – 50 × 10^6^ CFU and α_21_ from -50 – 1 × 10^6^ CFU into 201 bins each. We have varied the bounds and the lattice sizes of the grids and there was no change in the qualitative results (Fig. S5). In this case, the Shannon’s entropy (16, 17), 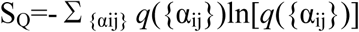 was maximized subject to the constraint that the estimated q({α_ij_}) should reproduce 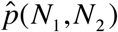, *i.e*.,

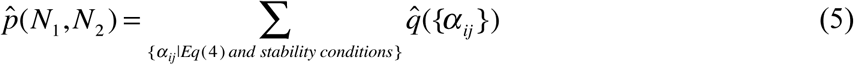

The stability conditions (Section 3, Supplementary Material) make sure that the fixed points in Eq. (4) are stable solutions. Eq. (5) was inverted to obtain 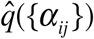. The solution is given by (14),

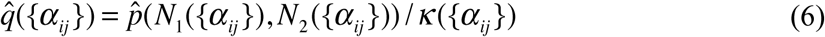

κ({α_ij_}) in the above equations denotes the degeneracy factor or the number of distinct points in the α space that produce the same value of N_1_ and N_2_. We calculated κ({α_ij_}) numerically by counting the number of lattice points in the α space that map to the same lattice point in the N-space.

### C. Determination of the role (*x*=*weak* or *x*=*strong*) of the mutant strain in co-infection kinetics

Our framework to evaluate the role of the mutant strain is divided into two main steps as outlined in the main text. <u>*Step 1*</u>. We estimated 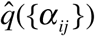 using the reference data set *_n_D_r_* as described in section B. We used 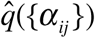 to generate models that belong in the *weak* or the *strong* category. The details regarding how these models were generated are given below. <u>*Weak models*</u>: A specific mutant strain is hypothesized to possess loss/gain of phenotype(s) at the design stage of the co-infection experiments (see Table I for details). The hypothesized changes in the specific phenotypes for the mutant strain will result in changes in a subset of LV interaction parameters pertaining to the co-infection kinetics of the populations of the mutant strain and another bacterial species within the chinchilla host. This subset of LV interaction parameters is denoted by {α_pq_}⊂ {α_ij_}, where p=i and q=j for each α_ij_ in {α_pq_}. *E.g*., for the co-infection with NTHi(*wt*)+Mcat(*mcaB*), the *mcaB* mutant strain increases the value of α_21_, thus the set {α_pq_} contains only one parameter α_21_; whereas for co-infection with NTHi(wt)+Mcat(*dtgt*), the *dtgt* mutant strain increases both α_21_ and α_22_, and {α_pq_} will contain two parameters, α_21_ and α_22_. The models in the *weak* category are defined by the probability distribution function of the interactions parameters (*q_w_*(*α*;**a**)) in the models. The *weak* models are parameterized by {a_k_} which quantifies the extent by which each of the interaction parameters in {α_pq_} is perturbed in 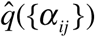 to generate *q_w_*(*α*;**a**), *i.e*.,

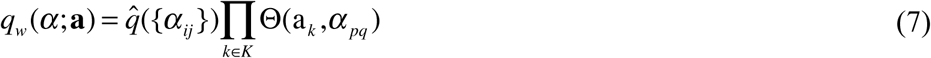

where, K ={(p,q)}. Θ(a_k_,α_pq_) denotes the Heaviside Theta function Θ(a_k_-α_pq_) or Θ(α_pq_-a_k_). Θ(a_k_-α_pq_) is chosen when the mutation decreases the value of α_pq_ such that the values α_pq_>a_k_ are absent in the mutant, or, Θ(α_pq_-a_k_) is chosen when the mutation increases the values of α_pq_ such that the lower values α_pq_<a_k_ are absent in the mutant. To illustrate, the weak model for the co-infection NTHi(*wt*)+Mcat(*mcaB*) is parameterized by a_1_ and is generated using 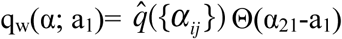. <u>*Strong models*</u>: The models in the *strong* category considered unanticipated changes in the LV interaction parameters for co-infection kinetics where a wild type strain is replaced by a mutant strain. If a mutant strain in a co-infection is hypothesized to change a subset of LV interaction parameters {α_pq_}, then the strong models consider changes in {α_pq_} and additional LV interactions (or unanticipated changes) outside of {α_pq_}. Similar to the weak models, the strong models are defined by the probability distribution function of the interactions parameters (*q_s_*(*α*;**a**)) and are parameterized by {a_k_} which quantifies the extent by which each of the interaction parameters is perturbed in 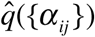 to generate *q*_s_(α;**a**), *i.e*.,

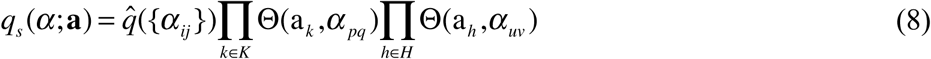

In the above equations, H ⊂ {α_ij_}\K, where H defines the set of LV interactions outside {α_pq_}. For example, the *strong* models for co-infection NTHi(*wt*)+Mcat(*mcaB*) were generated from in total 6 types of models: 3 types that vary pairs of interactions simultaneously, namely, 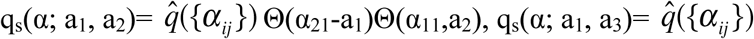 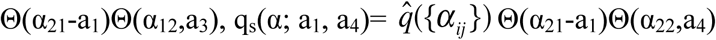, and, 3 types that vary a triplet of interactions simultaneously, namely, 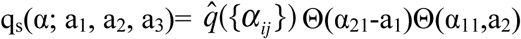 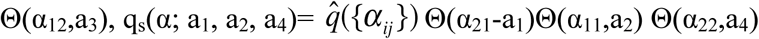, and, q_s_(α; a_1_, a_3_, a_4_)= 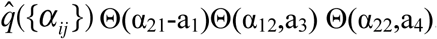. Note, in each of the six types of models α_21_ is always included and varied in the same way (increasing α_21_). We consider changes up to the triplet of LV interactions for generating the *strong* models.

<u>*Step 2*</u>. The models *q_w_*(*α*;**a**) and *q_s_*(*α*;**a**) were used to generate means of bacterial populations 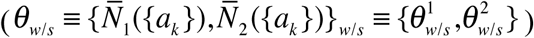 in the hosts using the steady state equations (Eq. (1)). We assumed that each of the *t* samples of the test dataset 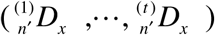 is distributed as a bivariate normal distribution,

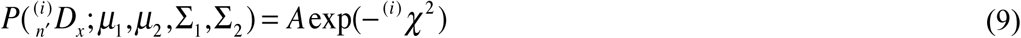

where,

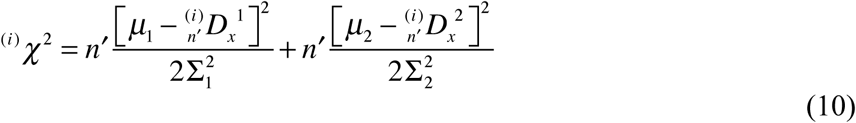

μ_1,2_ and Σ_1,2_ can be estimated from 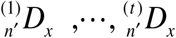 as,

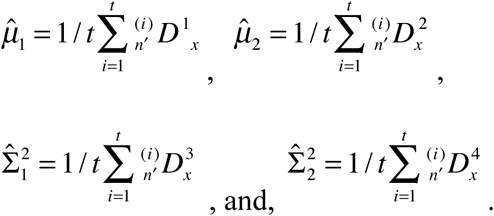

Note the *j* index in the superscript of 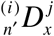 denotes the *j*^th^ element of the set 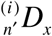 and should not be confused as a power. We demand that the means of the bacterial populations in the models describe the means in 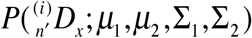, i.e.,

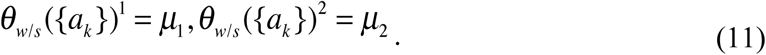

Next we determined the parameters ({a_k_}) by minimizing ^(i)^χ^2^ (or maximizing the corresponding likelihood),

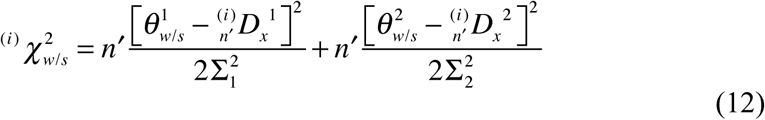

and then computed AIC for the i*^th^* sample of the test dataset after the minimization as,

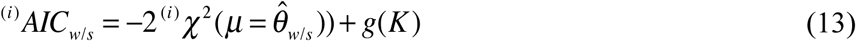

where, 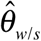 is the mean bacterial populations in the model evaluated at the {a_k_} values that minimized Eq. (10) and, K is the number of parameters in {a_k_}. We used *g(K)=2K* for our calculations.

#### Evaluation of the Condorcet winner

We made head-to-head comparisons for all possible pairwise combinations of models using AIC values as the metric. For a given pair, the winning model was the one which had a lower AIC value for a majority of the t samples. The model which won all of its head-to-head comparisons was declared the overall winner; this model is also known as the Condorcet winner, because it is preferred more than all others in pairwise comparisons. Throughout our study, both for the *in silico* and *in vivo* portions, we always found a Condorcet winner. The category under which the Condorcet winning model fell (*weak* or *strong*) was then assigned to x.

### D. Generation of the synthetic data

The purpose of the synthetic data was to evaluate the framework with data that mimics *in vivo* data but has known levels of mutation strength and host immune response. In addition to the reference dataset, we generated thirty-six total *in silico* mutation datasets. Specifically, we generated a mutation on each of the four parameters (α_11_, α_12_, α_21_, α_22_); a mutation had one of three levels of severity (low, moderate, large), and was paired with one of three levels of host’s immune response (none, weak, strong). Below, we outline how each was generated.

#### The Reference Set _n_D_r_

We first created a reference dataset analogous to a NTHi (*wt*)+Mcat (*wt*) co-infection experiment. Using the simplified LV two-species model (Eqn 4) we set the following ranges for each of the interaction parameters: α_11_ ϵ [2.74×10^-3^, 0.2], α_12_ ϵ [-200, 5], α_21_ ϵ [-5, 0.1], and α_22_ ϵ [1.89189, 140]. These ranges were guided by the values observed from the *in vivo* data so that the *in silico* reference set was roughly similar to *in vivo* data. To calculate a single data point of a (*N_1_, N_2_*) pair of abundances, we randomly drew a value for each parameter (assuming a uniform distribution) and calculated the steady state abundances using Eqn 4. For all the datasets, we generated n=10^5^ points per dataset. The reference dataset *_n_D_r_* was calculated by collating all the (N_1_, N_2_) pairs obtained by drawing the α parameters from random distributions as described above.

#### Mutation Sets

All “mutations” of the reference dataset were done by increasing the value of exactly one interaction parameter; specifically, by raising the minimum value in the parameter’s range. The three levels of the mutation indicated the severity of the increase. A low level α_11_ mutation, has α_11_ ϵ [0.027, 0.2]; at moderate severity: α_11_ ϵ [0.1, 0.2], and at large severity: α_11_ ϵ [0.18, 0.2]. The ranges for all twelve possibilities are shown in Fig. S2 in the supplement.

We also aimed to evaluate the framework with respect to different levels of a host’s immune response. In cases with no host response, we use the above method. For a non-zero immune response we used a simplified model wherein only *N_2_* induces the immune response, and only *N_1_* is affected (detrimentally). Specifically, we introduced a Michaelis-Menten term to the first ODE in the standard LV two-species model:

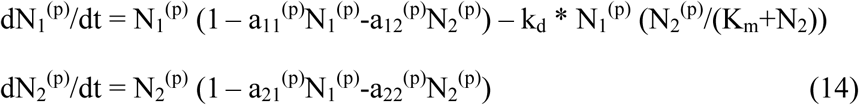

In that term, K_m_ is the amount of N_2_ necessary for the immune response to be at half-max. The susceptibility of N_1_ to an immune attack is governed by the k_d_ parameter. So, with no immune response, we set k_d_ = 0. For a weak immune response k_d_ ϵ [0, 1]. For a strong immune response, k_d_ ϵ [0, 10]. As before, when choosing a value for k_d_, we assumed a uniform distribution.

## Data Availability

The *in vivo* datasets and the MATLAB code used to analyze the data are available at https://datadryad.org/review?doi=doi:10.5061/dryad.j89d064

## Funding

This work was supported by grants (R01GM103612) from NIGMS to JD and (RO1DC10051) to WES.

## Author Contribution

VL, WS, WES, and JD planned research. LT and WES carried out experiments. VL, WS, SM, JD carried out in silico simulations and mathematical calculations, and analyzed in silico results. VL, WES, JD analyzed experimental data. VL, WS, WES, and JD wrote the manuscript.

## Research Ethics

Not required.

## Animal Ethics

All chinchilla infections were performed according to protocols approved by the Wake Forest Animal Care and Use Committee. This study was conducted according to the guidelines outlined by National Science Foundation Animal Welfare Requirements and the Public Health Service Policy on the Humane Care and Use of Laboratory Animals. The Institutional Animal Care and Use Committees (IACUC) at Wake Forest School of Medicine approved these animal studies. Wake Forest’s IACUC oversees the welfare, well-being and proper care and use of all vertebrate animals used for research and educational purposes at Wake Forest School of Medicine. The approved protocol number for the project is A13-140 and the Wake Forest School of Medicine animal welfare Assurance Number is D16-00248.

## Permission to carry out fieldwork

Not applicable.

## Acknowledgements

We thank Ali Snedden at the High Performance Computing (HPC) Center at the Research Institute at the Nationwide Children’s Hospital for technical help with the computation. The computation time provided by the Baker cluster at HPC is also acknowledged. We thank Veronica Vieland for a critical reading of the manuscript.

## Competing Interests

The authors declare no competing interests.

## 1. Basic statistics of *in vivo* bacterial abundance data

Table S1 shows some basic statistical analysis of all the *in vivo* data collected. The statistics include the mean, variance and covariance of the bacterial populations. *In vivo* data was collected at two time points: 7 and 14 days post inoculation. In addition to the wild type strains, various mutant strains were studied.

**Table S1.**
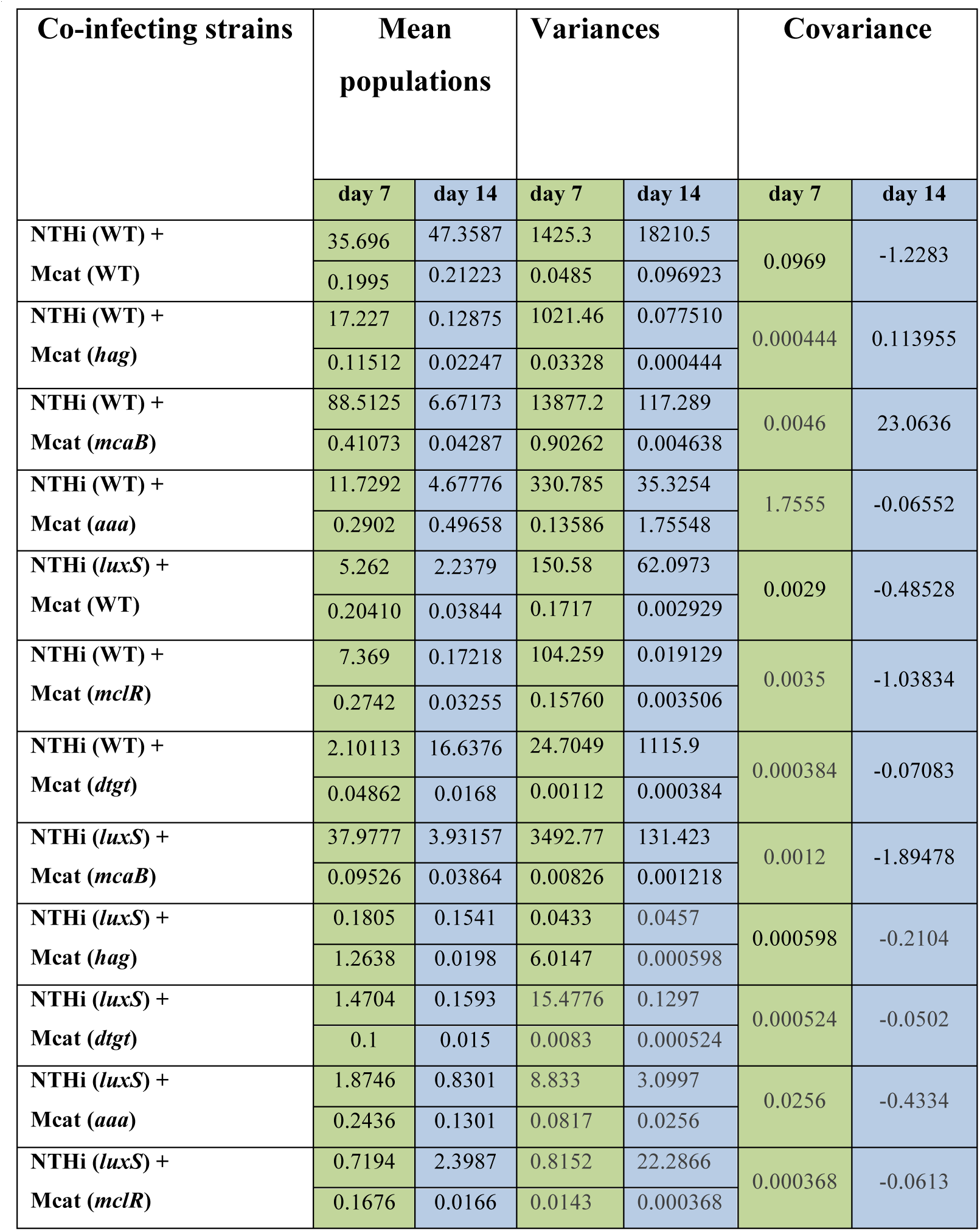
Means, variances and covariances for the populations of NTHi and Mcat strains measured in chinchilla middle ears.

## 2. Details of the MaxEnt calculation for estimation of P(N_1_,N_2_) using the data for the wild type strains

We used a MaxEnt approach to model the NTHi(WT) + Mcat(WT) co-infection data. Specifically, the P(N_1_,N_2_) model was in the form

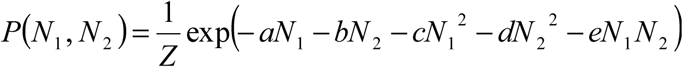

*Z* is the normalization parameter. The remaining parameters were calculated such that the means of N_1_ (*a*) and N_2_ (*b*), variances of N_1_ (*c*) and N_2_ (*d*), and the covariance (*e*) matched the measured data. These five parameters in the exponent were numerically solved for using the ‘fsolve’ function in Matlab. We tried 100,000 random initial guesses for the parameters and chose the set which minimized the error in each of the five statistical measures mentioned above.

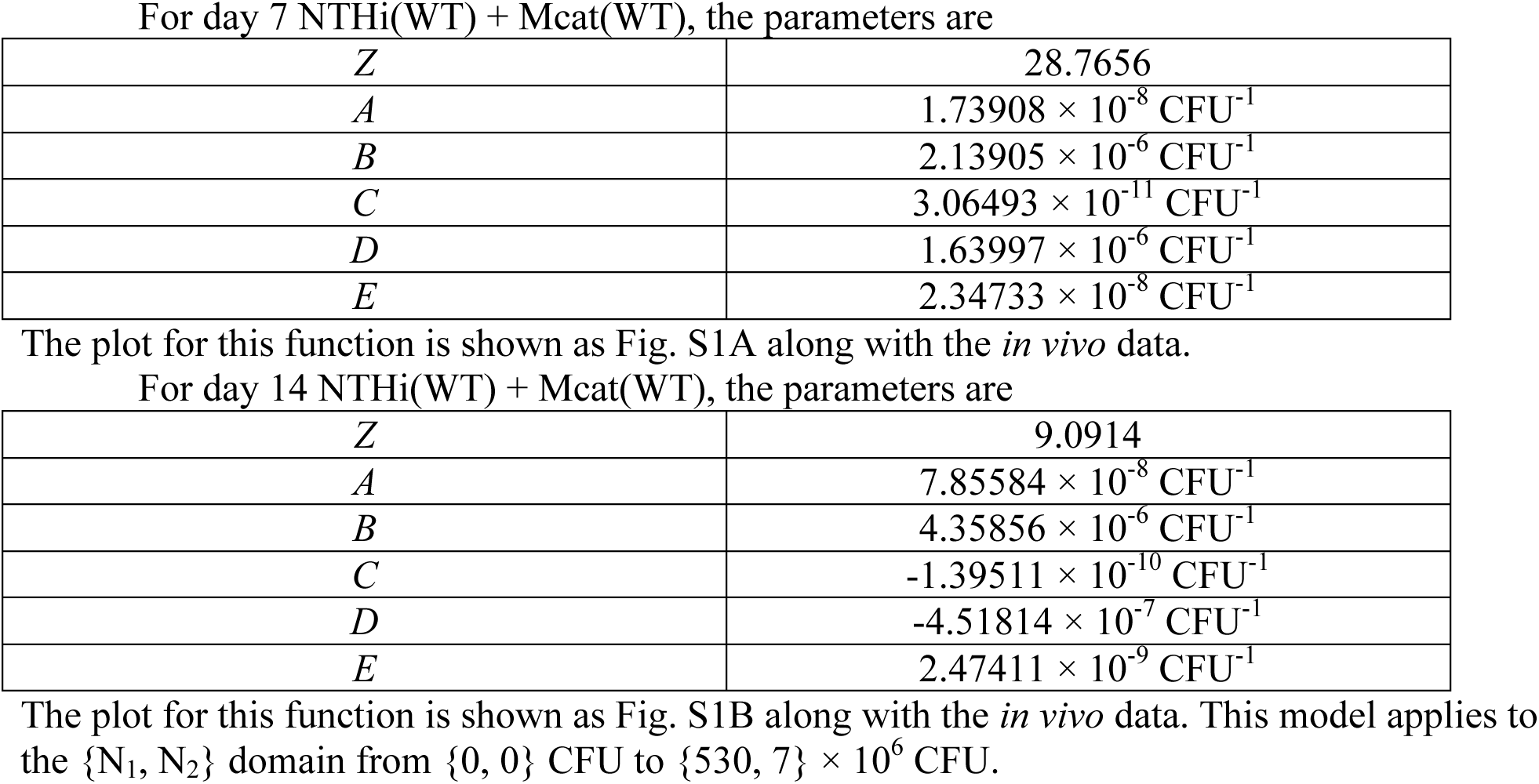

|  |  |
| --- | --- |
| $Z$ | 28.7656 |
| $A$ | $1.73908 \times 10^{-8} \text{ CFU}^{-1}$ |
| $B$ | $2.13905 \times 10^{-6} \text{ CFU}^{-1}$ |
| $C$ | $3.06493 \times 10^{-11} \text{ CFU}^{-1}$ |
| $D$ | $1.63997 \times 10^{-6} \text{ CFU}^{-1}$ |
| $E$ | $2.34733 \times 10^{-8} \text{ CFU}^{-1}$ |
| $Z$ | 9.0914 |
| $A$ | $7.85584 \times 10^{-8} \text{ CFU}^{-1}$ |
| $B$ | $4.35856 \times 10^{-6} \text{ CFU}^{-1}$ |
| $C$ | $-1.39511 \times 10^{-10} \text{ CFU}^{-1}$ |
| $D$ | $-4.51814 \times 10^{-7} \text{ CFU}^{-1}$ |
| $E$ | $2.47411 \times 10^{-9} \text{ CFU}^{-1}$ |

**Figure S1.**
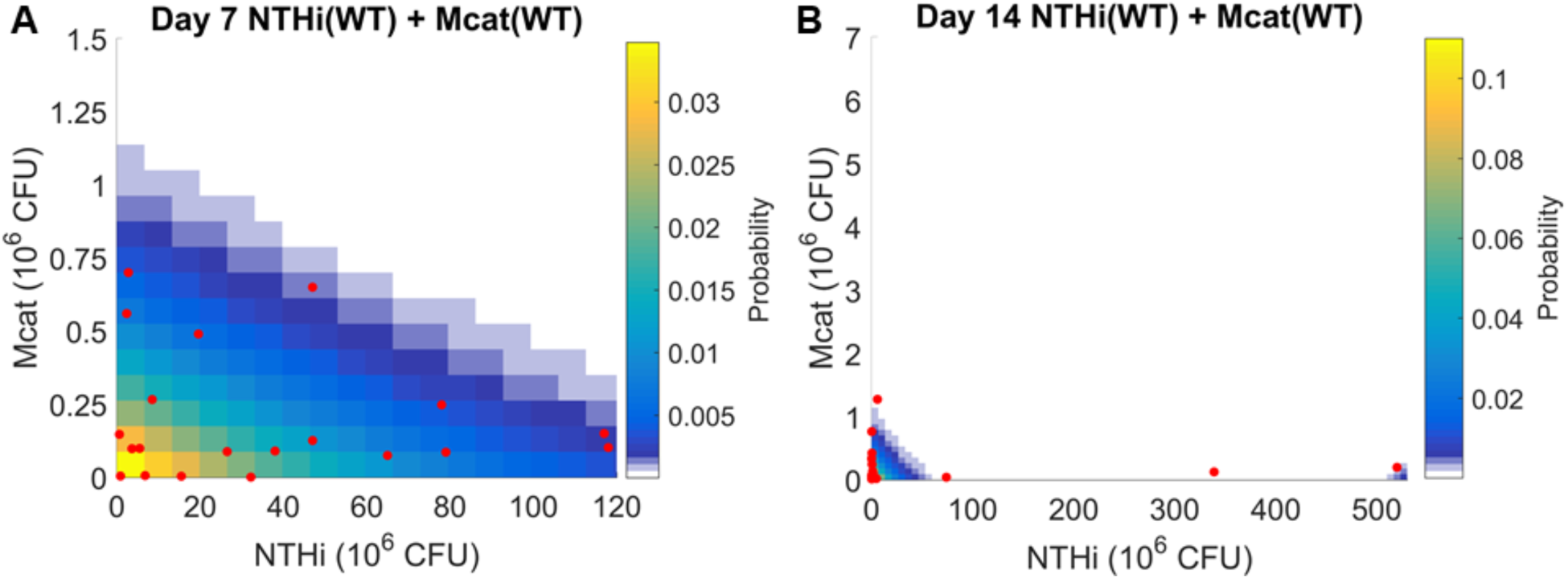
Host-host variations of NTHi (wt) and Mcat (wt) populations in co-infection experiments. **(A)** Shows populations of NTHi and Mcat in the counts (red points) obtained from the bullae of two ears of each animal for 10 chinchillas at day 7 post inoculation. The heat map shows the probability distribution function 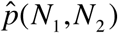 estimated by our MaxEnt based method. The axes have been zoomed into the region containing all *in vivo* data. **(B)** Measurements for day 14 post inoculation shown using the same visualization scheme as in (A). The heat map shows the MaxEnt estimation for 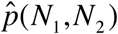 corresponding to the data at day 14.

## 3. Derivation of Stability Analysis

Starting with Eq. (3), we have the following two equations describing the abundance of both species at steady state:

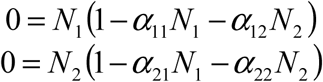

The stability matrix, *S*, which describes the derivative of each species with respect to all others near the steady state can be constructed. In general, for *n* total species, an element of *S* is given by

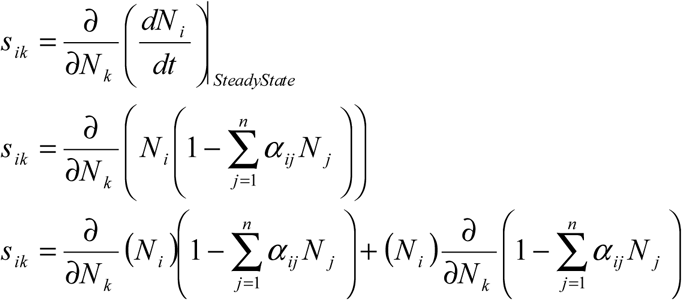

The first term is *0*, because 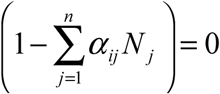 at Steady State.

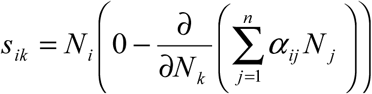

The second term is *0* for all *j*, except when *j*=*k*:

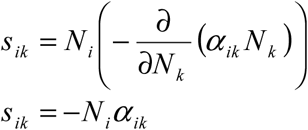

For our two species system, the stability matrix can be written as

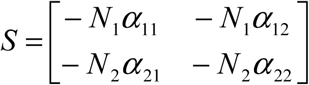

We then calculate the Eigen values of *S*. The steady state is considered stable if and only if the Real parts of every Eigen value is less than or equal to 0.

## 4. Two parameter marginal distributions of Q(α_11_, α_12_, α_21_, α_22_)

From the P(N_1_,N_2_) fit of the reference data, we use a MaxEnt method to calculate Q(α_11_, α_12_, α_21_, α_22_). These four parameters correspond to the LV parameters (Fig. 1). α_11_ (>0) and α_22_ (>0) represent intra- species interactions for NTHi and Mcat, respectively. α_12_ and α_21_ represent the effect of Mcat on the growth of NTHi, and, NTHi on the growth of Mcat, respectively. α_12_ and α_21_ can be positive (competitive interaction), zero (neutral interaction), or, negative (cooperative interaction) based on Eq. (3). We calculate the six pairwise correlations and plot the marginal distributions to visualize these correlations. Fig. S2 shows the Q(**α**) for day 7, and Fig. S3 shows the Q(**α**) for day 14. These correlations between the α parameters have contributions from three sources: (i) the interdependence between them via Eq. (3), (ii) shape of the distribution of P(N_1_,N_2_) and (iii) the stability criterion (Supplemental Section 3) that ensures that N_1_ (>0) and N_2_ (>0) in Eq. (3) are stable fixed points of the kinetics in Eq. (1).

**Figure S2.**
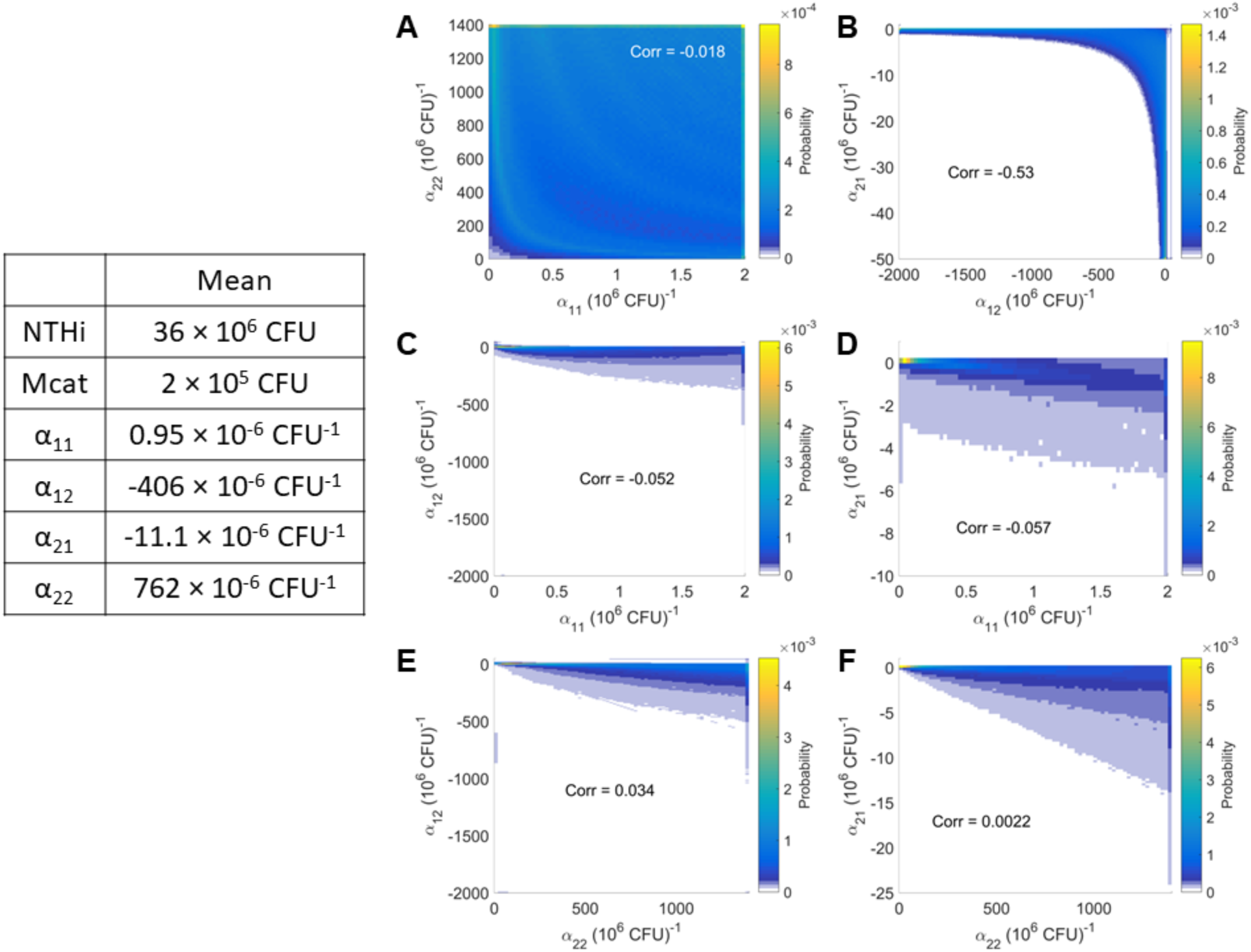
Day 7 Estimation of LV interactions in the co-infection kinects with wild type strains. The two dimensional joint distributions of the interactions, Q(α_11_, α_12_, α_21_, α_22_), are shown in six planes: **(A)** α_11_-α_22_, **(B)** α_12_-α_21_, **(C)** α_11_-α_12_, **(D)** α_11_-α_21_, **(E)** α_22_-α_12_, and, **(F)** α_22_-α_21_ using heat-maps. The table shows the mean values of the populations of NTHi and Mcat and the interaction parameters. The Person correlations (Corr) between the interaction parameters are shown on the figures.

**Figure S3.**
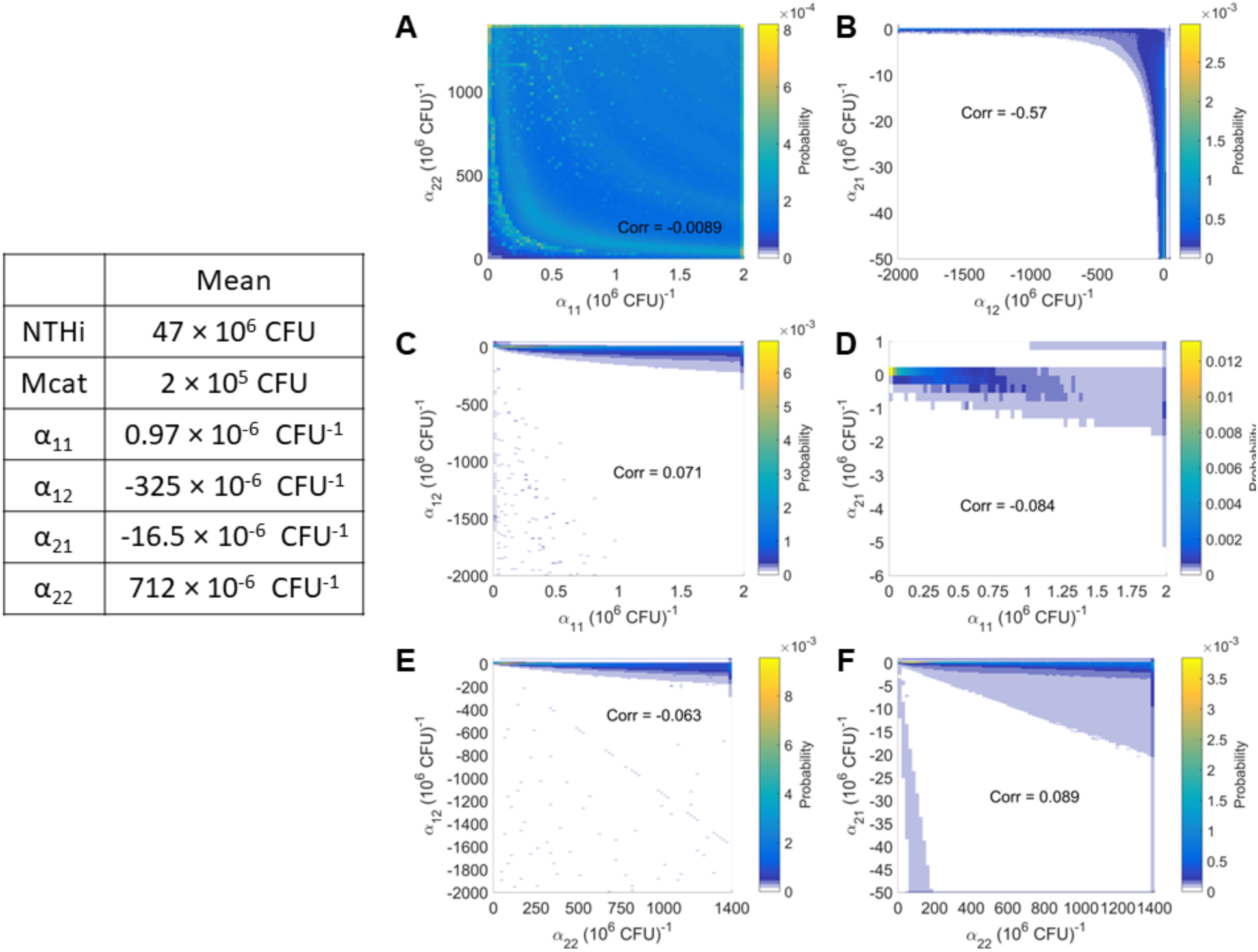
Day 14 Estimation of LV interactions in the co-infection kinects with wild type strains. The two dimensional joint distributions of the interactions, Q(α_11_, α_12_, α_21_, α_22_), are shown in six planes: **(A)** α_11_-α_22_, **(B)** α_12_-α_21_, **(C)** α_11_-α_12_, **(D)** α_11_-α_21_, **(E)** α_22_-α_12_, and, **(F)** α_22_-α_21_ using heat-maps. The table shows the mean values of the populations of NTHi and Mcat and the interaction parameters. The Person correlations (Corr) between the interaction parameters are shown on the figures. This analysis is done using the same domains in NTHi-Mcat and **α**-space as the day 7 data.

## 5. Comparison between the Condorcet winner model and measurement at day 14 post inoculation with mutant strains

In the main text we present the results for bacterial kinetics as observed 7 days post co-infection. These results are presented with a model (called the Condorcet winner) which best describes the changes in the mutation strain data compared to wild type. We were also able to analyze data from 14 days post inoculation (Fig. S4).

**Figure S4.**
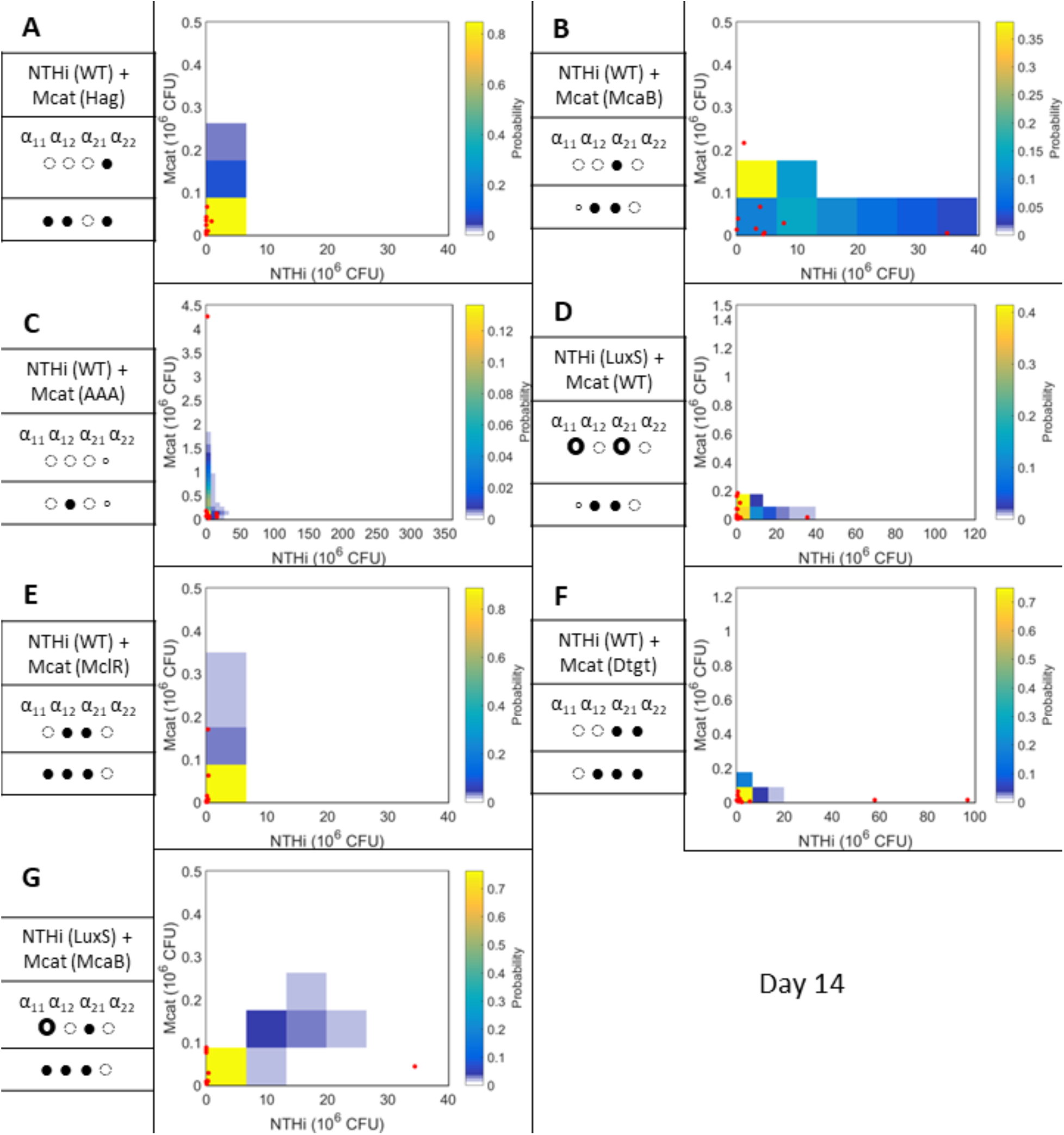
Day 14 post inoculation: data and predicted models. The data are displayed using the same visualization scheme as Fig. 4 in the main text. The following combinations are shown: **(A)** NTHi(WT)-Mcat(*hag*). **(B)** NTHi(WT)-Mcat(*mcaB*). **(C)** NTHi(WT)-Mcat(*aaa*). **(D)** NTHi(*luxS*)-Mcat(*aaa*). **(E)** NTHi(WT)-Mcat(*mclR*). **(F)** NTHi(WT)-Mcat(*dtgt*). **(G)** NTHi(*luxS*)-Mcat(*mcaB*). All the mutant strains studied show unanticipated changes in the interactions *in vivo* that *strongly* regulate bacterial kinetics. The specific regulations are indicated by the Condorcet winning model. Fig. 4 shows at day 7, some strains’ unanticipated changes are *weak* regulators of kinetics. This result indicates that the host immune response plays a more significant role in bacterial kinetics by day 14 than day 7.

## 6. Doubling the range of the α-domain does not change the results

As stated in the main text, we chose a fixed domain in α-space; that is, α_11_ ϵ [0.027, 2], α_12_ ϵ [-2000, 50], α_21_ ϵ [-50, 1] and α_22_ ϵ [18.9189, 1400]. To verify that this chosen domain is sufficiently large and the results are independent of this choice, we repeated our analysis with a domain roughly twice as large (Fig. S5). That is, α_11_ ϵ [0.0541, 4], α_12_ ϵ [-4000, 100], α_21_ ϵ [-100, 5] and α_22_ ϵ [37.8378, 2800]. The lattice size was proportionally increased with the domain.

**Figure S5.**
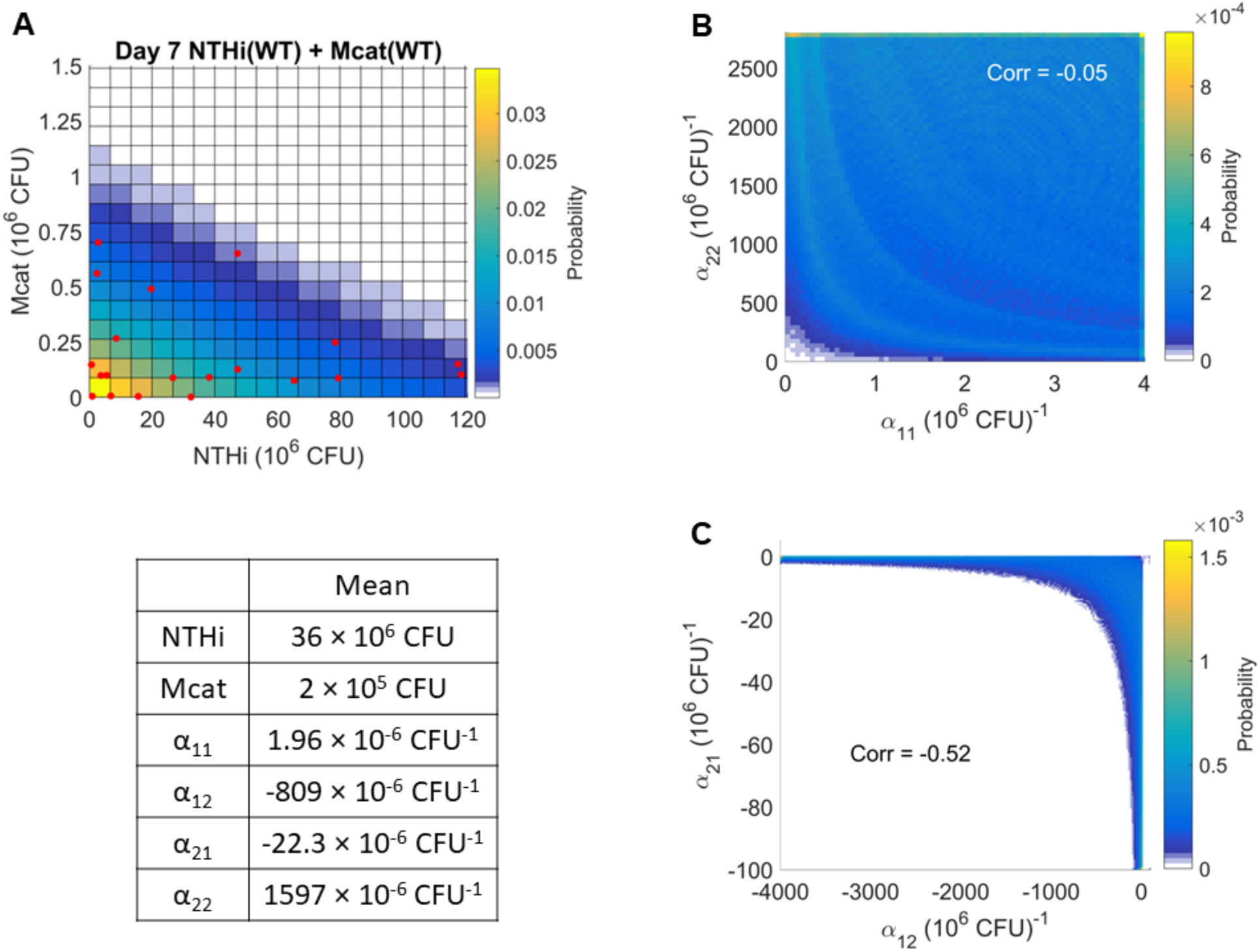
Doubled α-domain. **(A)** Shows the NTHi and Mcat populations at day 7 post inoculation along with the MaxEnt estimated P(N_1_,N_2_). This panel is a reproduction of Fig. S1A. **(B)** and **(C)** show Q(α_11_, α_22_) and Q(α_12_, α_21_) respectively. These panels are analogous to Figs. S2A – S2B. Note that the correlations are similar, and because the domains were doubled in size, the average α-values are also doubled. Using this larger α-domain, we recalculated the predictions for the day 7 NTHi(WT) + Mcat(Hag) and day 7 NTHi(WT) + Mcat(McaB) co-infection experiment. The resulting predictions (data not shown) were equivalent to the ones shown in Figs. 4A – 4B.

## 7. *In silico* Reference Data

We generated three sets of *in silico* reference data; each estimated a specific level of the host’s immune response. Below we show these three datasets (N_1_, N_2_) pairs, the 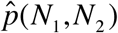 fits and the corresponding 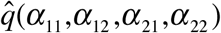.

**Figure S6.**
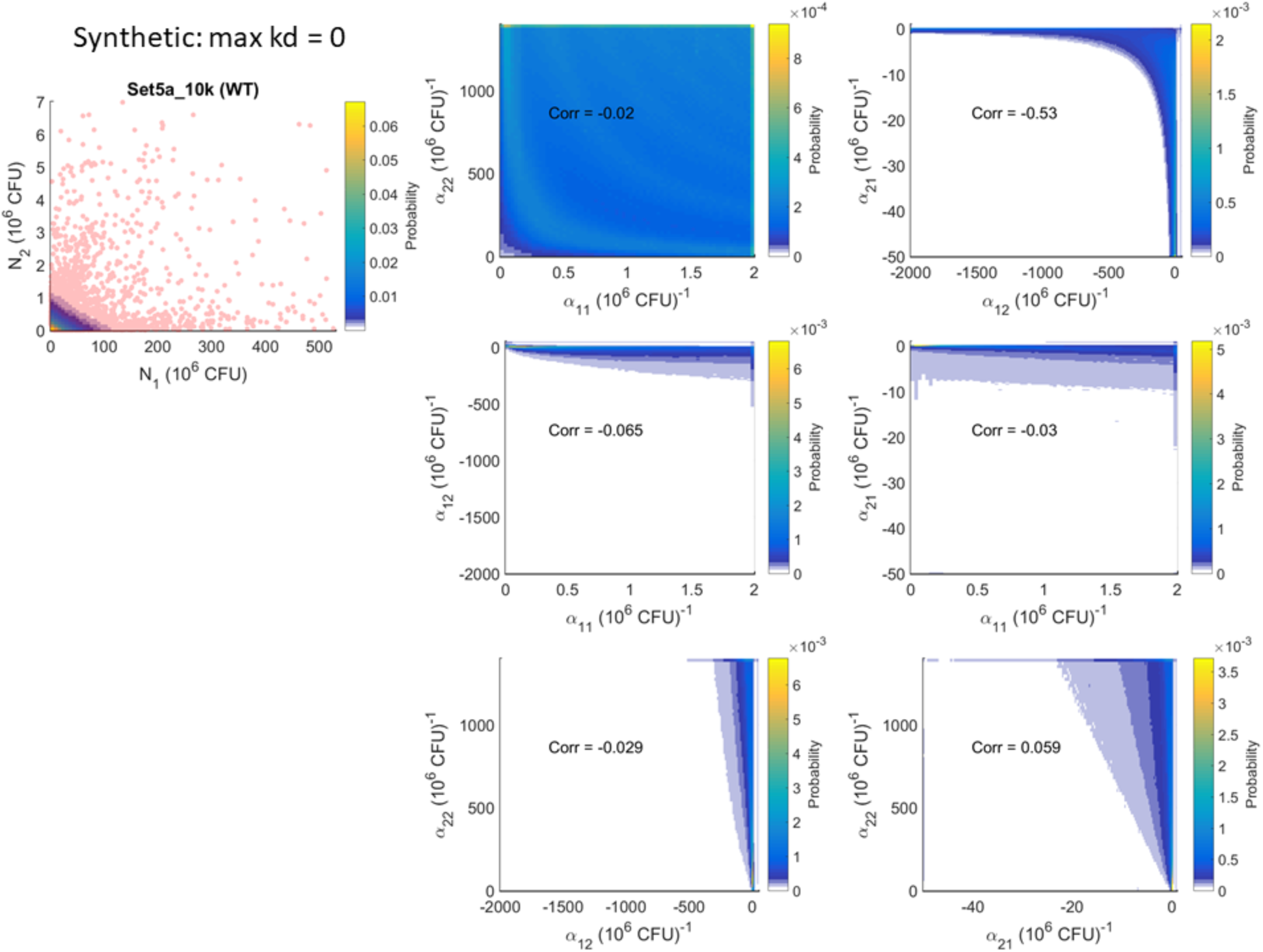
No Immune Term. In the absence of an immune response, we generated 10^4^ pairs of (N_1_, N_2_) shown here as the red dots. The fit, 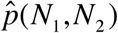, is shown underlying the scattered points quantified using the color bar. For all six pair-wise combinations of the 4 parameters (α_11_, α_12_, α_21_, α_22_), we show the marginal distributions as in Fig. S2.

**Figure S7.**
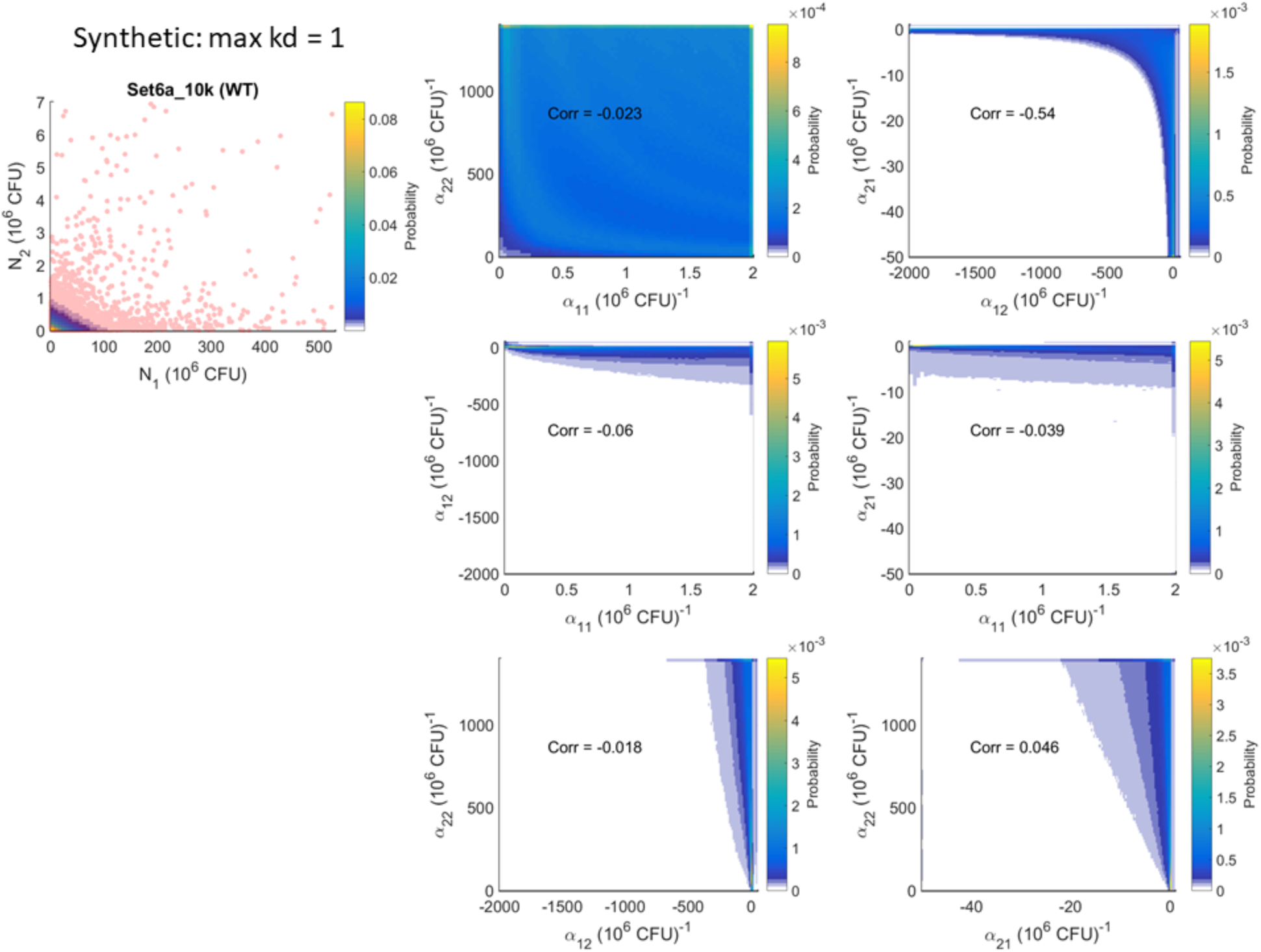
Weak Immune Term. To model a weak immune response, as described in the main text, we allow only N_2_ to solicit an immune response, and only N_1_ to be susceptible to inhibition by this response. This inhibition is weak (dictated by k_d_). The panels are similar to Fig. S6.

**Figure S8.**
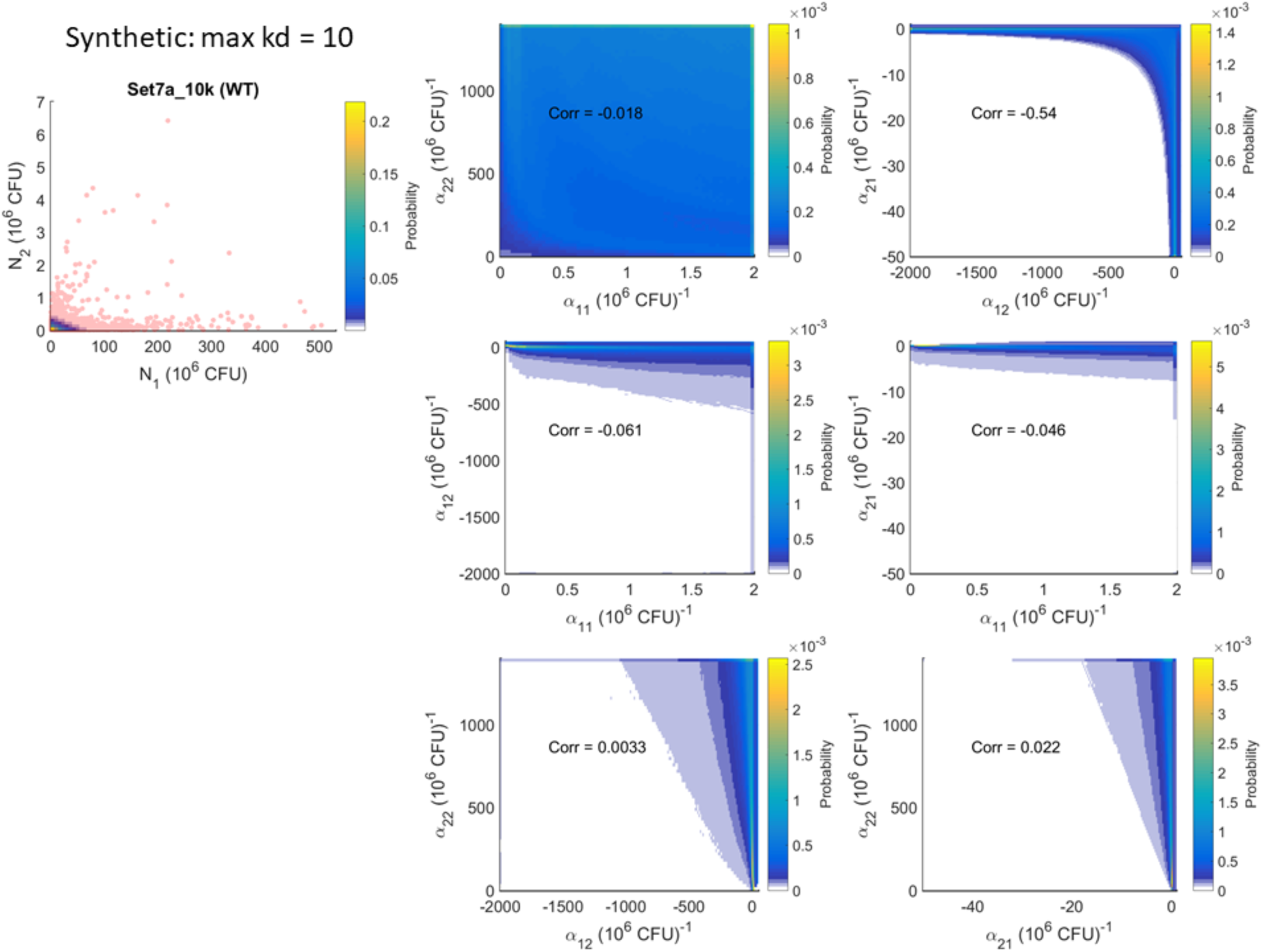
Strong Immune Term. Only N_2_ solicits an immune response, and only N_1_ is susceptible to inhibition by the immune response. This inhibition is strong (dictated by k_d_). The panels are similar to Fig. S6.

## 8. Summary of all *in silico* mutant strain results

We generated 36 datasets *in silico* based on three levels of the immune term, three levels of the severity of the mutation and four mutations (one for each LV parameter). We applied our framework to each of these datasets and determined if the unanticipated changes in the interactions were *weak* or *strong* regulators of the bacterial kinetics. If *strong*, then we went further and identified the specific nature of the unanticipated regulation. Figs. S9 – S12 show these results. The winning models are denoted by the changes in {α_11_, α_12_, α_21_, α_22_} for the wild-type+wild-type co-infection. O indicates no change, X indicates an increase, and, o indicates a decrease.

**Figure S9.**
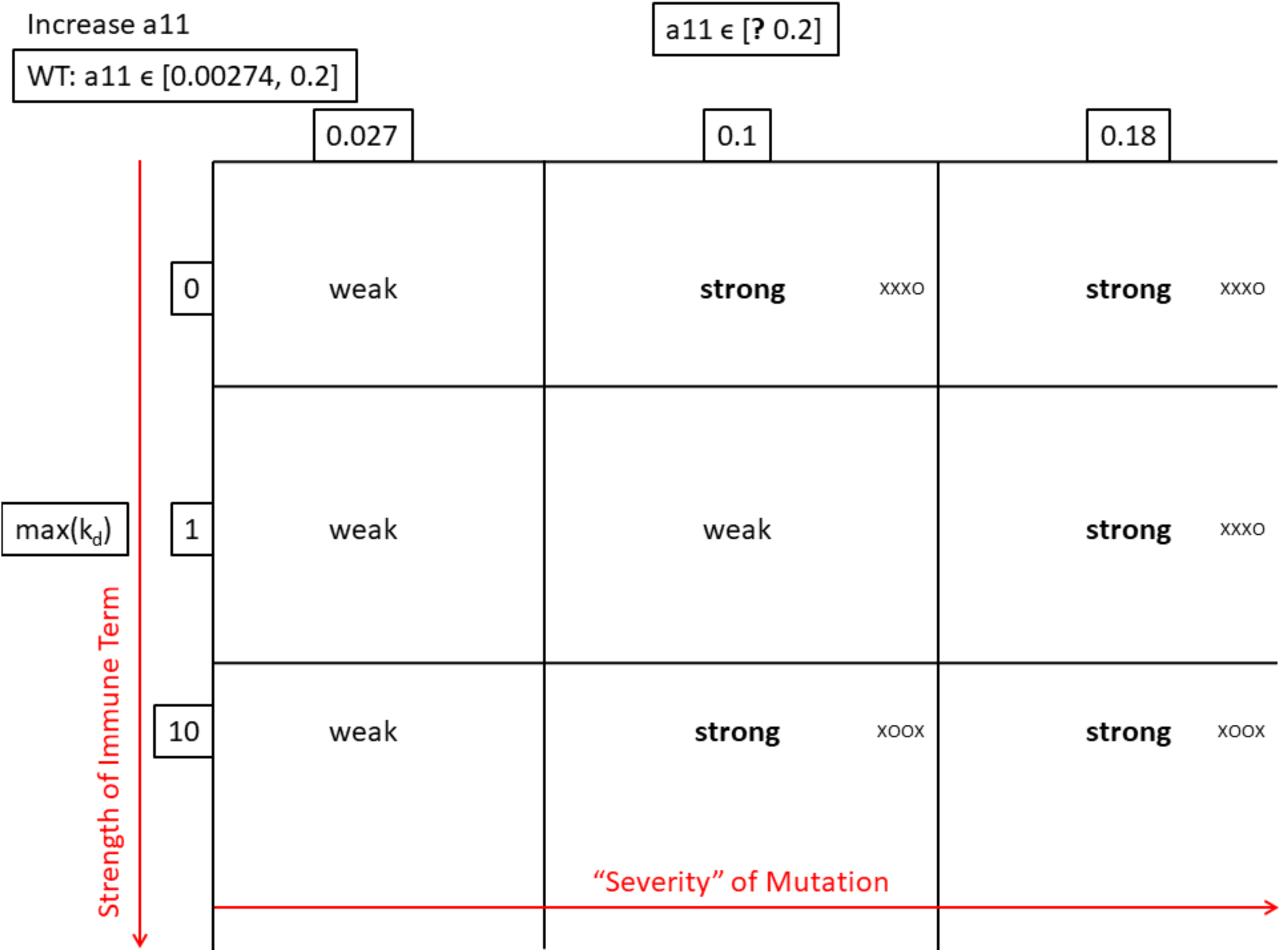
Mutation of α_11_. We varied the strength of the immune term (increasing by row from top to bottom) and the severity of the mutation (increasing by column from left to right). The immune term is dictated by the k_d_ parameter (see Eq. (14)); the minimum is 0 and the maximum is indicated at each row. The severity of the mutation is dictated by the range of α_11_; the wild type’s range is shown on the top left. The values above the columns indicate the new minimum for each case.

**Figure S10.**
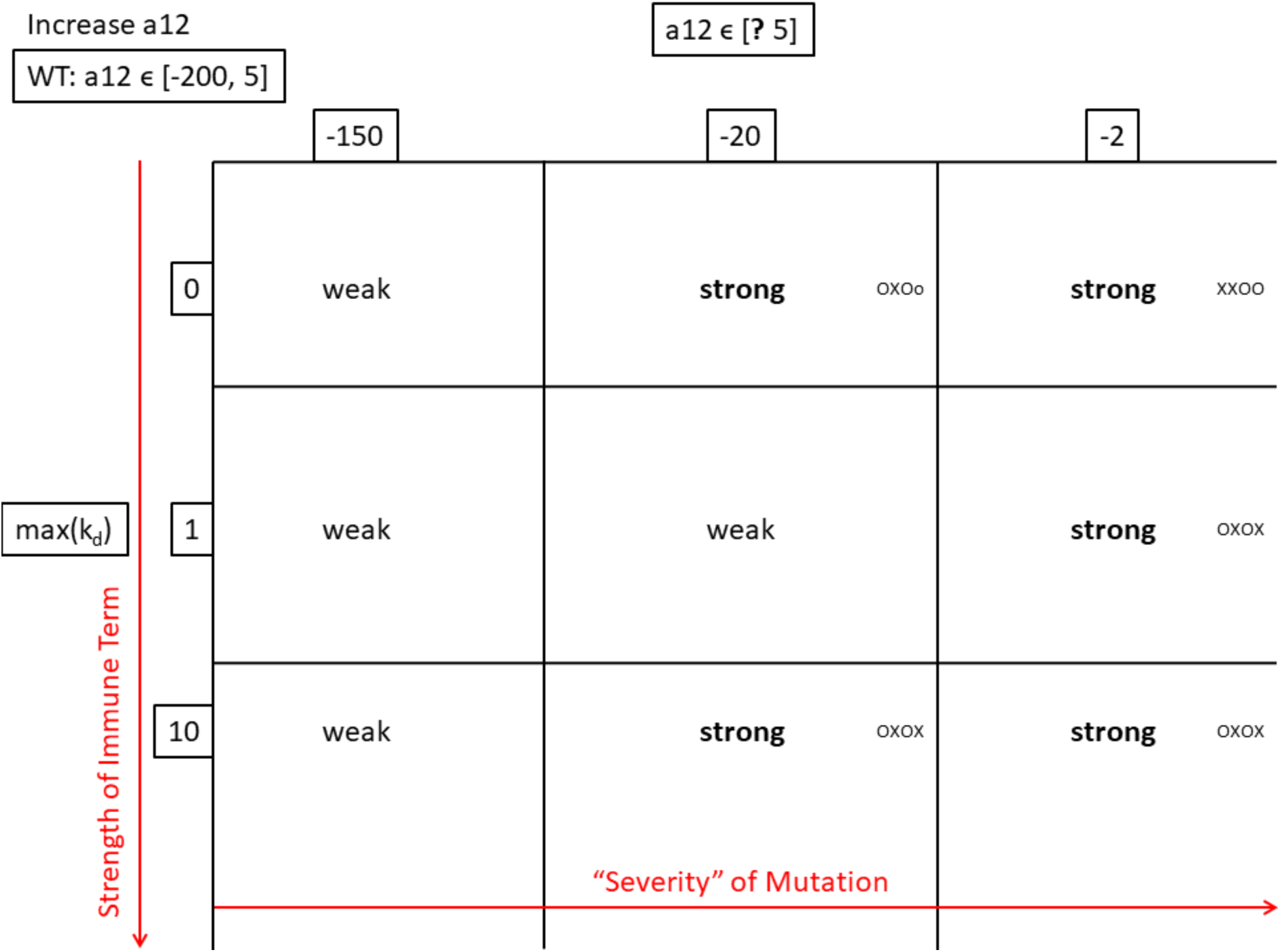
Mutation of α_12_. The figure is arranged similar to Fig. S9. The severity of the mutation is dictated by the range of α_12_; the wild type’s range is shown on the top left.

**Figure S11.**
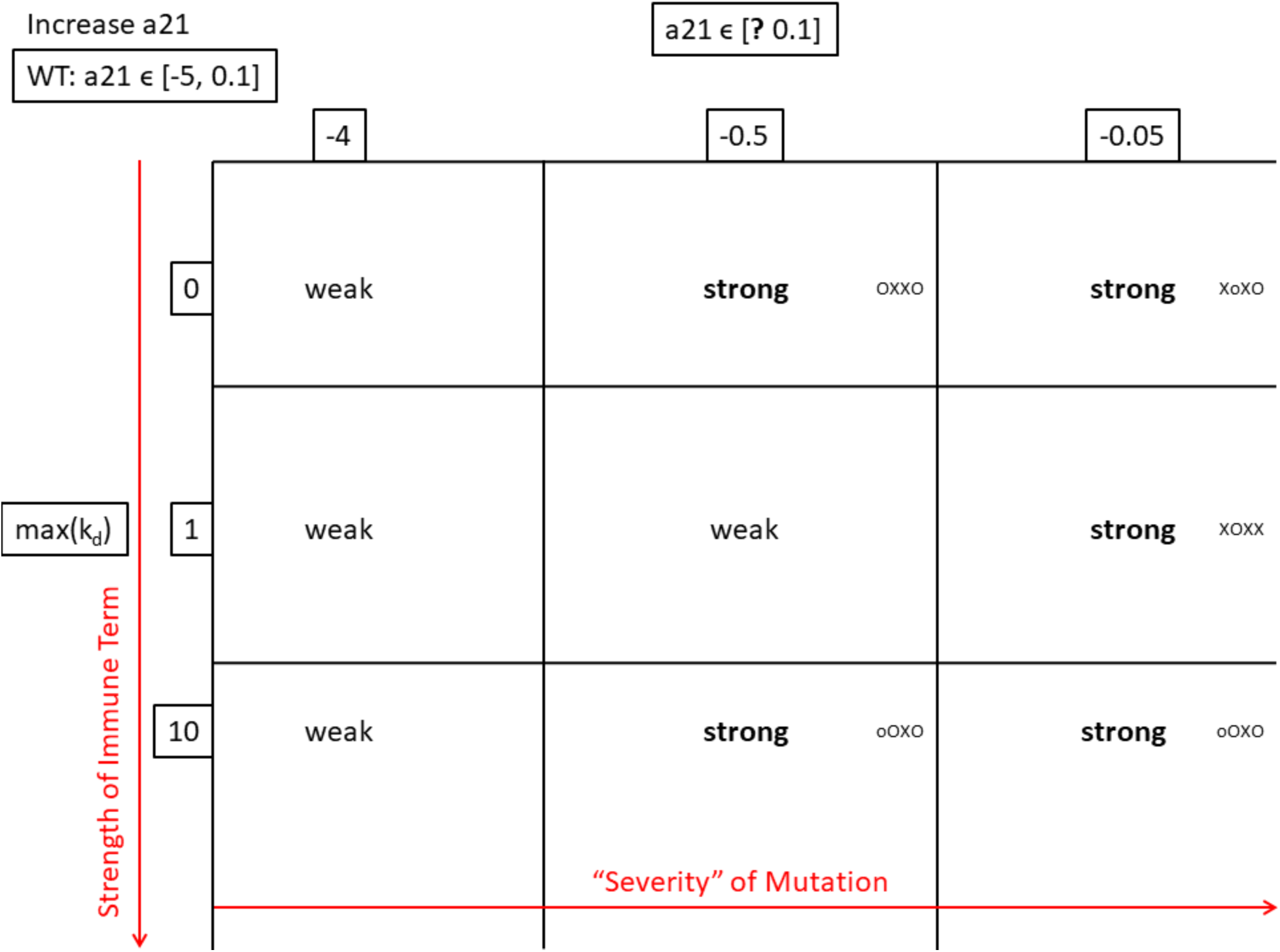
Mutation of α_21_. The figure is arranged similar to Fig. S9. The severity of the mutation is dictated by the range of α_21_; the wild type’s range is shown on the top left.

**Figure S12.**
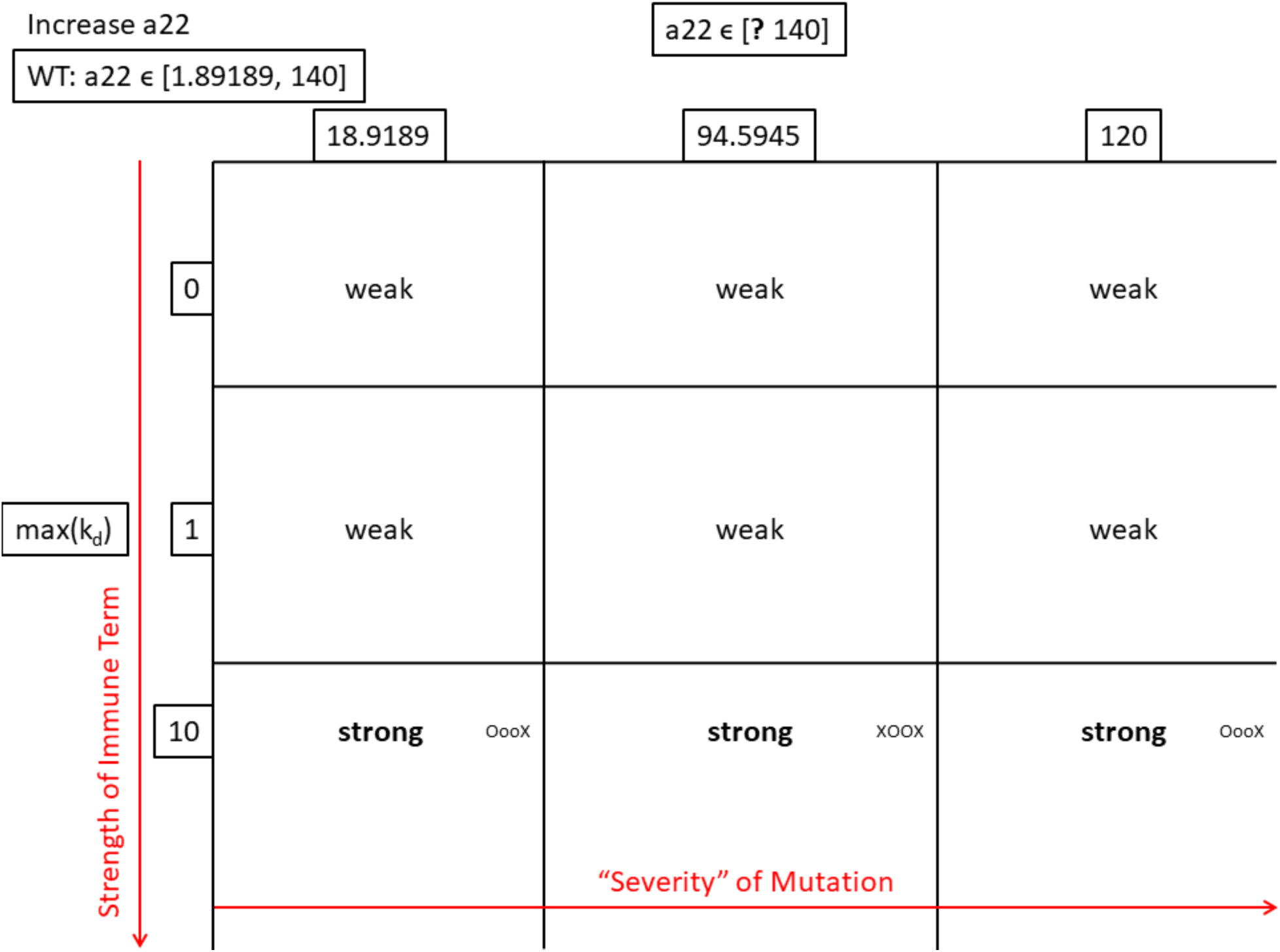
Mutation of α_22_. The figure is arranged similar to Fig. S9. The severity of the mutation is dictated by the range of α_22_; the wild type’s range is shown on the top left.

